## Supplementary for "High genome plasticity and frequent genetic exchange in *Leishmania tropica* isolates from Afghanistan, Iran and Syria"

**Table S1.** The number of total, valid, and discarded sequence reads per sample, along with the numbers of candidate and valid SNPs per sample. *Samples from the same patient. The association between the numbers of initial reads and candidate SNPs was high ($r^2$=0.243) compared to the association between the numbers of valid read and valid SNPs ($r^2$=0.028), suggesting that quality-control improved the accuracy of the true variation present.

| Name | Total | Post_trim | Discarded | Candidate_SNPs | Valid_genome_SNPs | Valid_chrom_SNPs | Valid_contig_SNPs |
|---|---|---|---|---|---|---|---|
| 07_01513 | 8,920,610 | 5,262,264 | 3,658,346 | 248,003 | 94,438 | 94,205 | 233 |
| 07_00242 | 16,802,189 | 9,622,500 | 7,179,689 | 231,544 | 94,283 | 94,037 | 246 |
| 13_01390 | 8,934,270 | 5,399,512 | 3,534,758 | 254,117 | 110,204 | 109,935 | 269 |
| 13_01024 | 4,630,516 | 4,353,757 | 276,759 | 286,877 | 142,232 | 141,909 | 323 |
| 13_01233 | 10,635,251 | 6,305,464 | 4,329,787 | 237,073 | 71,639 | 71,439 | 200 |
| 13_00550 | 5,726,620 | 4,234,418 | 1,492,202 | 232,728 | 69,843 | 69,662 | 181 |
| 14_00771 | 13,431,305 | 12,818,607 | 612,698 | 303,553 | 268,558 | 267,856 | 702 |
| 14_00849 | 6,728,629 | 6,728,629 | 0 | 296,291 | 200,596 | 200,124 | 472 |
| 14_01223 | 9,082,097 | 9,082,097 | 0 | 284,973 | 195,692 | 195,217 | 475 |
| 14_00642 | 20,605,192 | 12,233,464 | 8,371,728 | 251,558 | 115,049 | 114,776 | 273 |
| 15_00019 | 9,307,904 | 9,307,904 | 0 | 293,452 | 196,107 | 195,684 | 423 |
| 15_01088 | 7,037,807 | 6,474,880 | 562,927 | 286,463 | 150,578 | 150,246 | 332 |
| 15_02015 | 4,583,057 | 4,583,057 | 0 | 293,584 | 220,540 | 220,064 | 476 |
| 15_02480 | 4,143,879 | 4,143,879 | 0 | 296,138 | 191,212 | 190,748 | 464 |
| 15_02576 | 6,400,586 | 6,400,586 | 0 | 298,612 | 220,748 | 220,207 | 541 |
| 15_02597 | 10,475,771 | 9,939,940 | 535,831 | 307,110 | 278,095 | 277,195 | 900 |
| 15_01620 | 15,719,096 | 9,017,253 | 6,701,843 | 246,353 | 90,360 | 90,190 | 170 |
| 16_00075 | 5,282,999 | 5,282,999 | 0 | 298,281 | 229,345 | 228,720 | 625 |
| 16_00674 | 7,331,460 | 6,991,655 | 339,805 | 300,241 | 230,378 | 229,798 | 580 |
| 16_00964 | 5,244,524 | 4,974,688 | 269,836 | 274,934 | 150,050 | 149,552 | 498 |
| 16_14706 | 5,374,855 | 5,098,282 | 276,573 | 297,764 | 220,422 | 219,861 | 561 |
| 17_01604* | 5,742,571 | 5,323,561 | 419,010 | 295,922 | 203,653 | 203,141 | 512 |
| AVG | 8,733,690 | 6,980,882 | 1,752,809 | 278,616 | 170,183 | 169,753 | 430 |
| MIN | 4,143,879 | 4,143,879 | 0 | 232,137 | 69,843 | 69,662 | 170 |
| SD | 4,401,636 | 2,576,343 | 2,648,495 | 25,792 | 63,894 | 63,717 | 186 |

**Figure S1**. The read coverage (A), mapping quality (MQ) (B), reverse reads (C) and forward reads (D) (all on x-axes) varied across the 22 samples (coloured dashed lines) compared to the number of SNPs called (y-axes). These plots illustrate that the relative numbers of SNPs (and therefore relative genetic distances) called across samples was consistent.

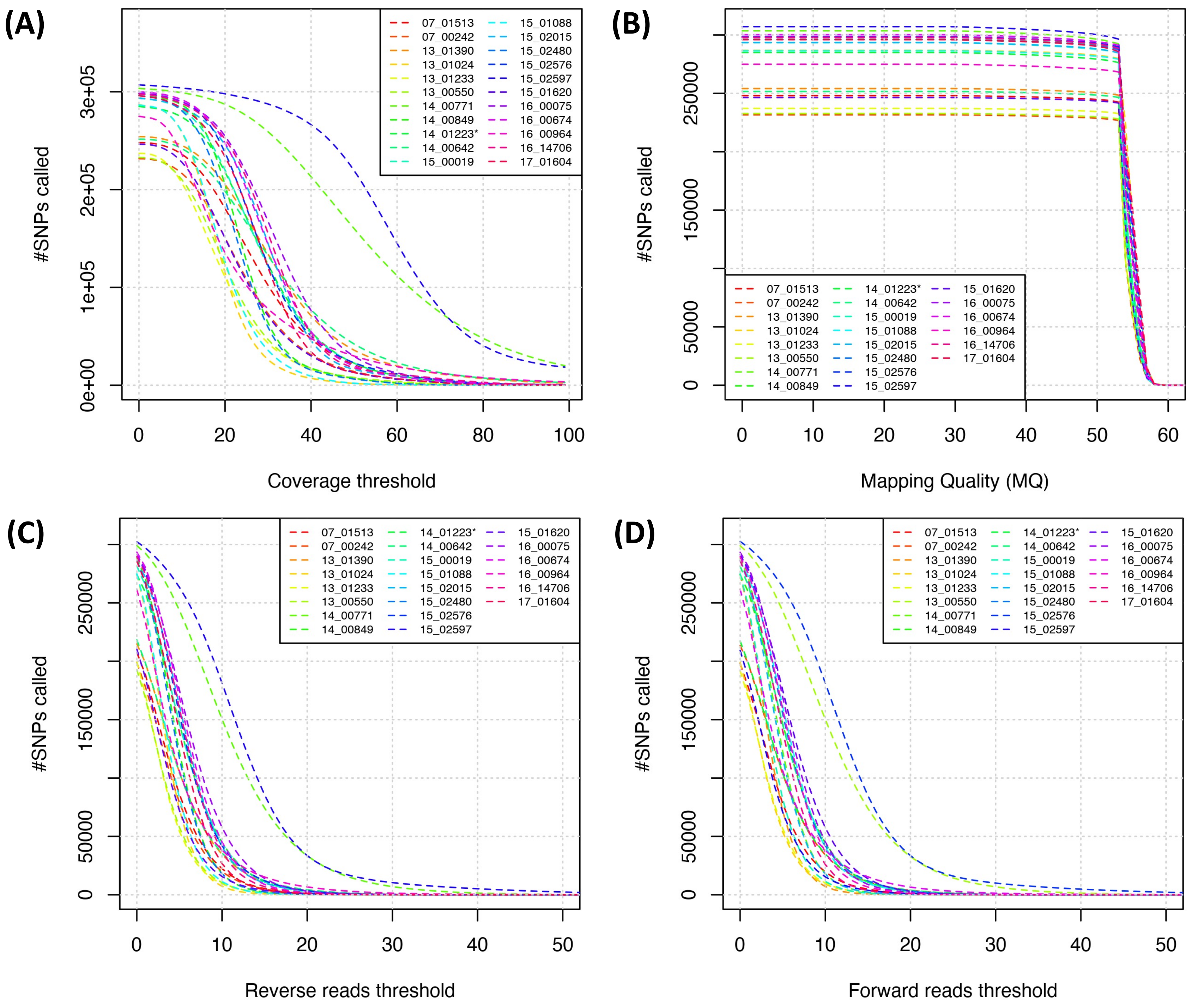

**Figure S2**. The numbers of homozygous (x-axis) and heterozygous (y-axis) chromosomal SNPs per sample. There was no association between the numbers of homozygous (x-axis) and heterozygous (y-axis) SNPs per sample (black circle). The 95% confidence interval for the adjusted $r^2$ of 0.07 (blue line) is in grey, showing a small negative correlation.

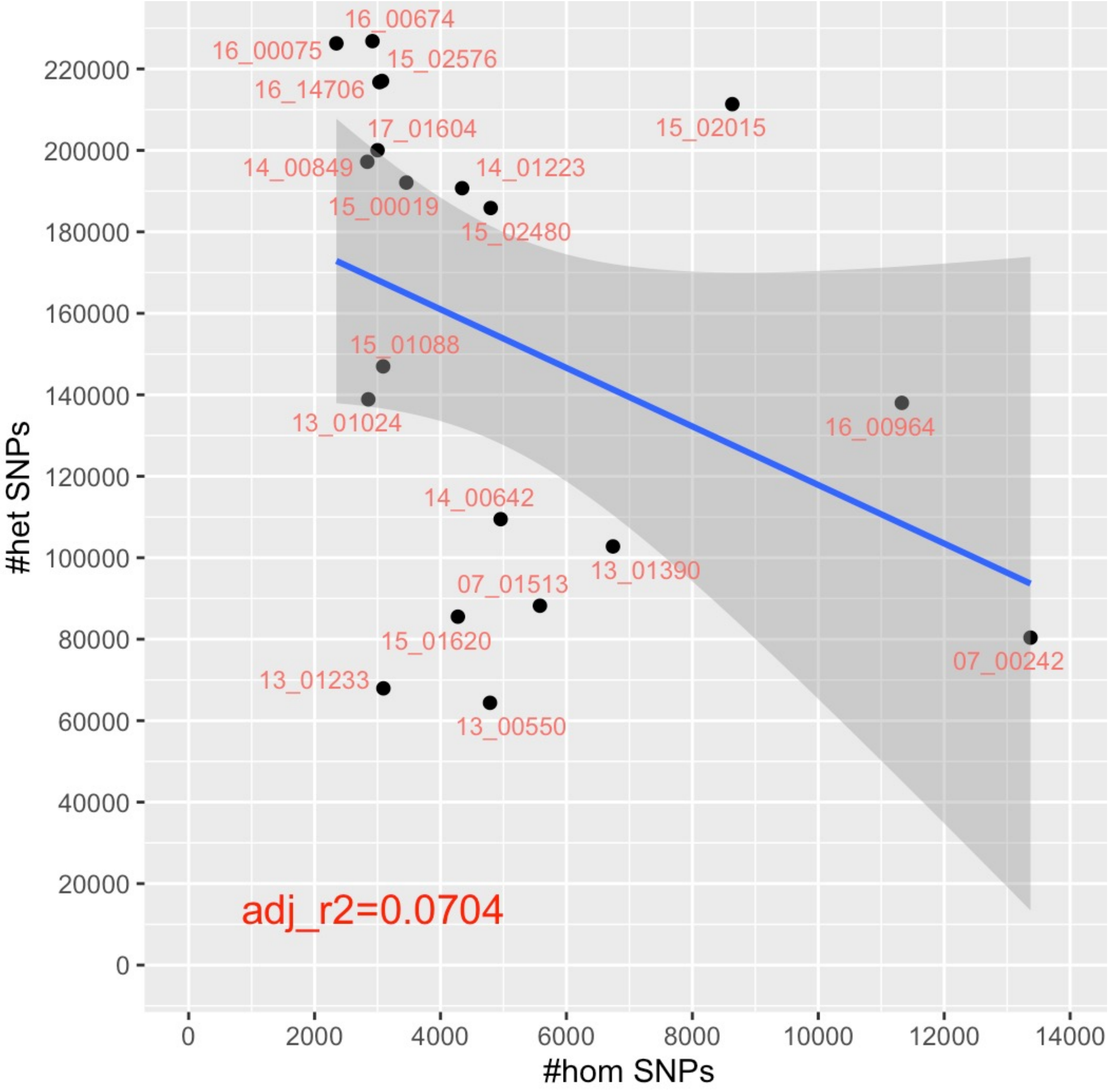

**Figure S3.** 14_01223 had evidence of near-homozygosity on chromosome 2 where it had 539 homozygous SNPs (>6 times more than all the other samples) and only 359 heterozygous SNPs (far fewer than the others). (A) This was illustrated by the SNPs' read-depth allele frequency (RDAF) distributions (top left) and the RDAF levels across the chromosome (top right). (B) A phylogeny constructed from chromosome 2's SNPs showing the relatedness of the 22 isolates with the *L. tropica* reference genome ("ref") and demonstrating that 14_01223 was genetically distinct. The inferred genetically distinct groups from FastBAPs are represented by the *L. tropica* reference (yellow area) and non-reference groups (purple area). (C) The homozygous (x-axis) and heterozygous (y-axis) SNPs per Kb for all 22 samples. Most samples including 14_01223 were approximately disomic for chromosome 2. There were no associations with *ori* regions or SSRs.

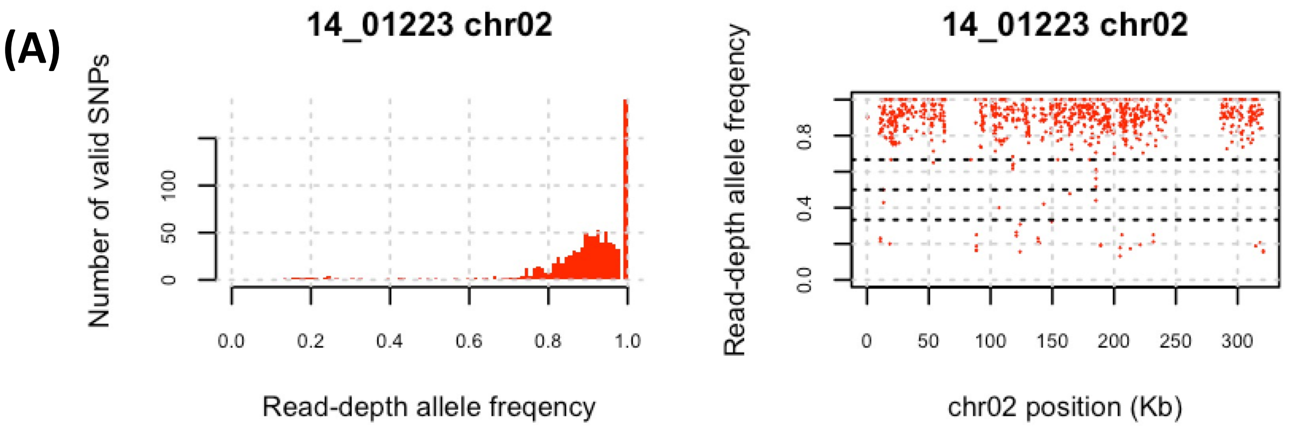


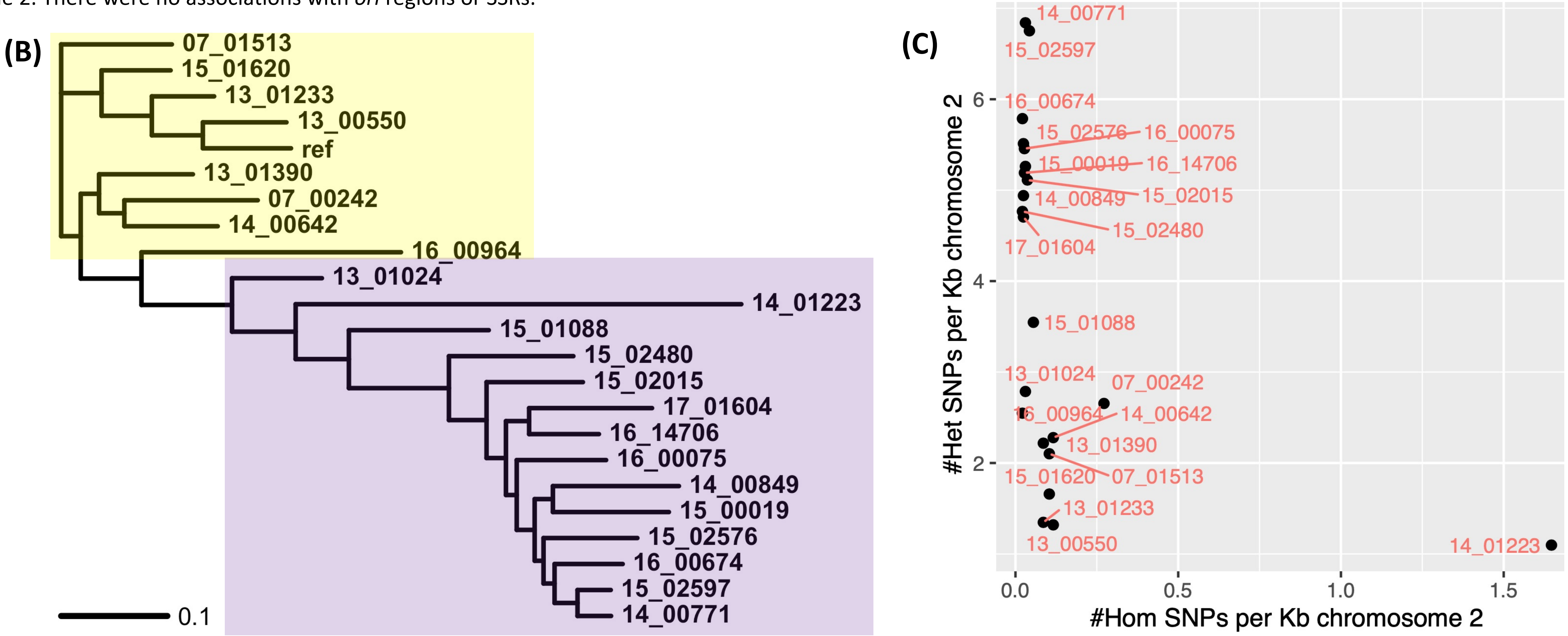

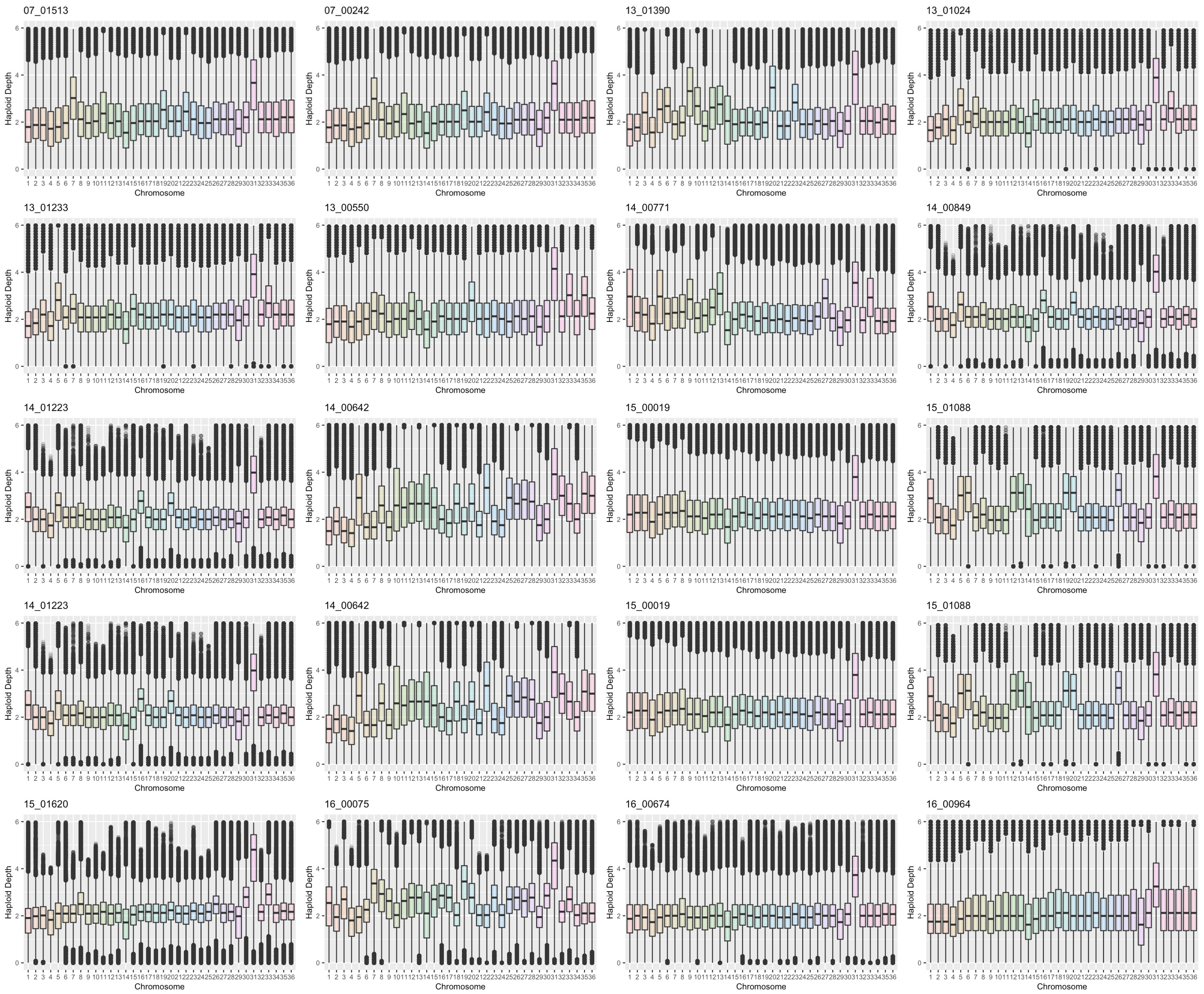


**Figure S4**. The chromosomes' (x-axis) distributions of their normalised haploid read depth per base (y-axis) for all 22 isolates. The box indicates the interquartile range with the median shown by the horizontal bar in the middle. Chromosomal regions with higher or lower normalised depth are shown by the whiskers and circles to indicate outliers. Most samples were mainly disomic, except for chromosome 31, which was mainly tetrasomic.

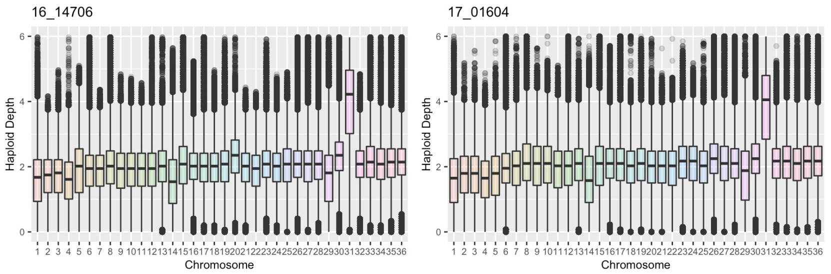

**Figure S5.** (A) Assignment of genetically distinct population using FastBAPS for SNPs at <1.28 Mb, 1.60-1.78 Mb, and > 1.80 Mb on chromosome 36 showing the genetic relatedness of the 22 isolates with the *L. tropica* reference genome ("ref", red group). All isolates were disomic, except 14_00642 was trisomic. (B) The homozygous (x-axis) and heterozygous (y-axis) SNPs per Kb for all 22 samples, highlighting the higher level of homozygous SNPs in 14_01223, 16_00075 and 16_00964. (C) 14_01223, 16_00075, 16_00964 and 07_00242 clustered in their own intermediate genetic group (red) due to unique variation 1.60-1.78 Mb, distinct from the non-reference group (yellow), compared to > 1.8 Mb (D).

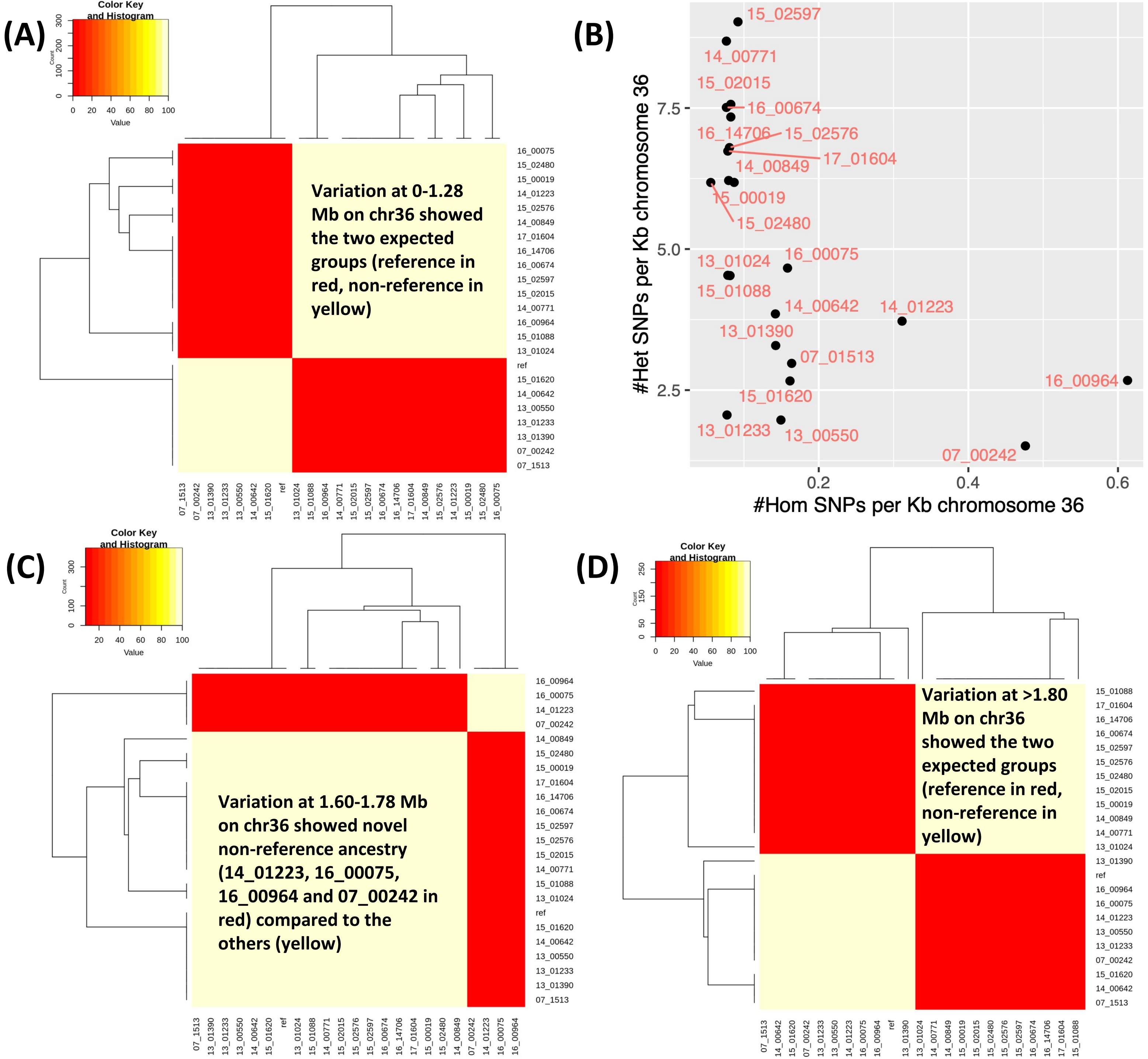

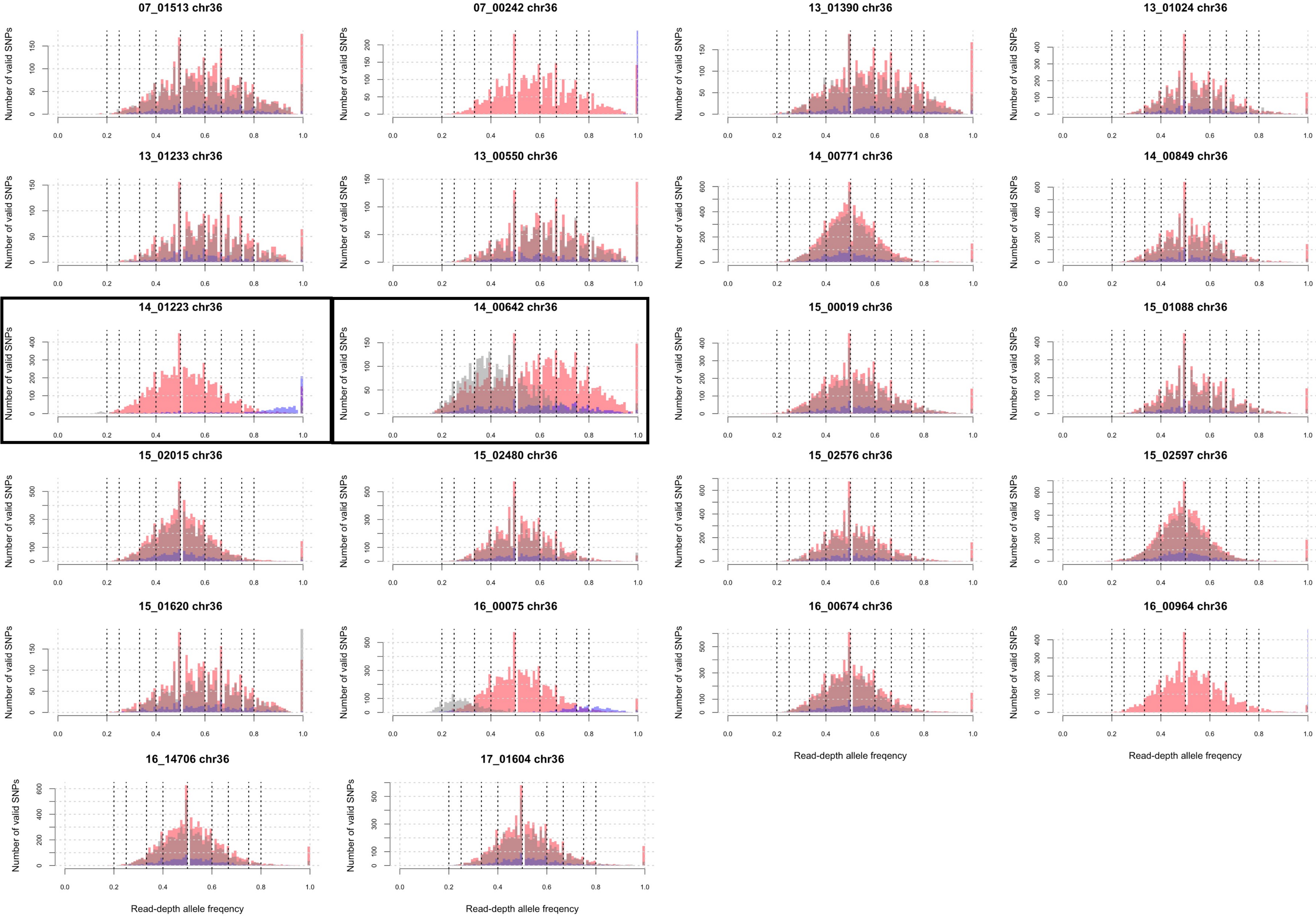


**Figure S6**. The read-depth allele (RDAF) distributions SNPs at chromosome 36's 5' end (<1280 Kb, red), middle (1600-1780 Kb, blue) and 3' end (>1800 Kb, grey). All isolates were disomic for chromosome 36, except 14_00642 was trisomic, which is reflected in its peaks at ~0.33 for the 3' end (grey) and at ~0.67 for the 5' end (red). 14_01223 (boxed) has heterozygosity for the middle region (red) and more homozygosity for the 5' end (blue). 16_00075 (boxed) had peaks at 0.75 for the 5' end, then 0.5 for the middle, and 0.25 for the 3' end.

**Figure S7**. The read-depth allele frequency (RDAF) levels for heterozygous SNPs (y-axis) across chromosomes (x-axis) shown as boxplots highlighting the interquartile range. Chromosome 23 had a lower RDAF rate (0.43±0.06) compared to the other chromosomes (average 0.52±0.04). Isolates with extreme RDAFs are named; these include those with lower than expected RDAFs: 14_01223 chr2, 07_00242 chr4, 07_01513 chr11, 15_02480 chr12, 07_00242 chr13, 16_00964 chr13, 07_00242 chr22, 14_00771 chr24, 13_01390 chr27, 15_02480 chr28, 16_00964 chr28, 16_00964 chr29, 07_00242 chr29, 07_00242 chr32, 15_02015 chr32, 07_00242 chr33, 14_00771 chr33, 16_00964 chr36, 07_00242 chr36; and those with higher than expected RDAFs: 14_00642 chr22, 07_00242 chr23, 13_01233 chr23, 15_02480 chr23, 16_00075 chr23, 16_00964 chr23.

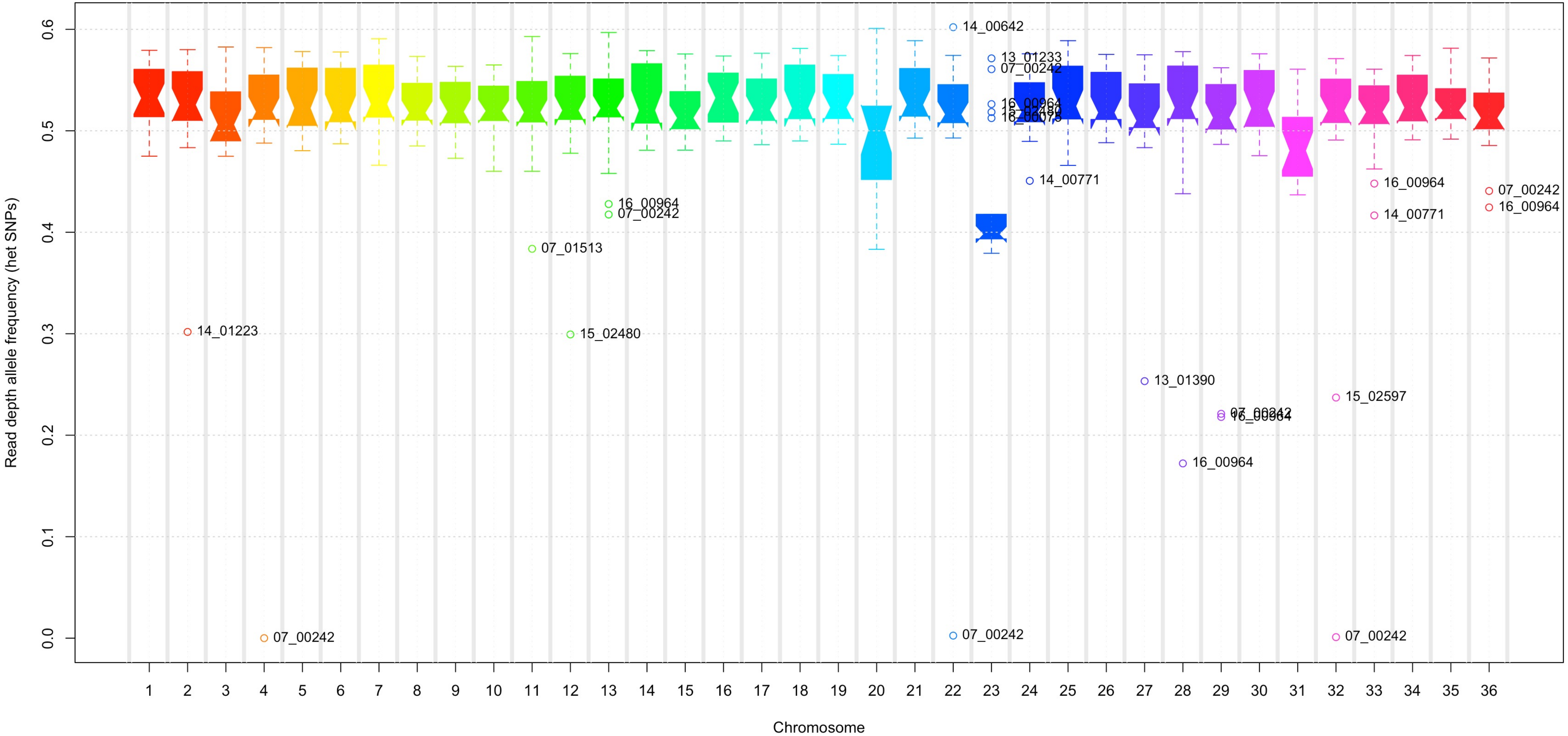

(A)

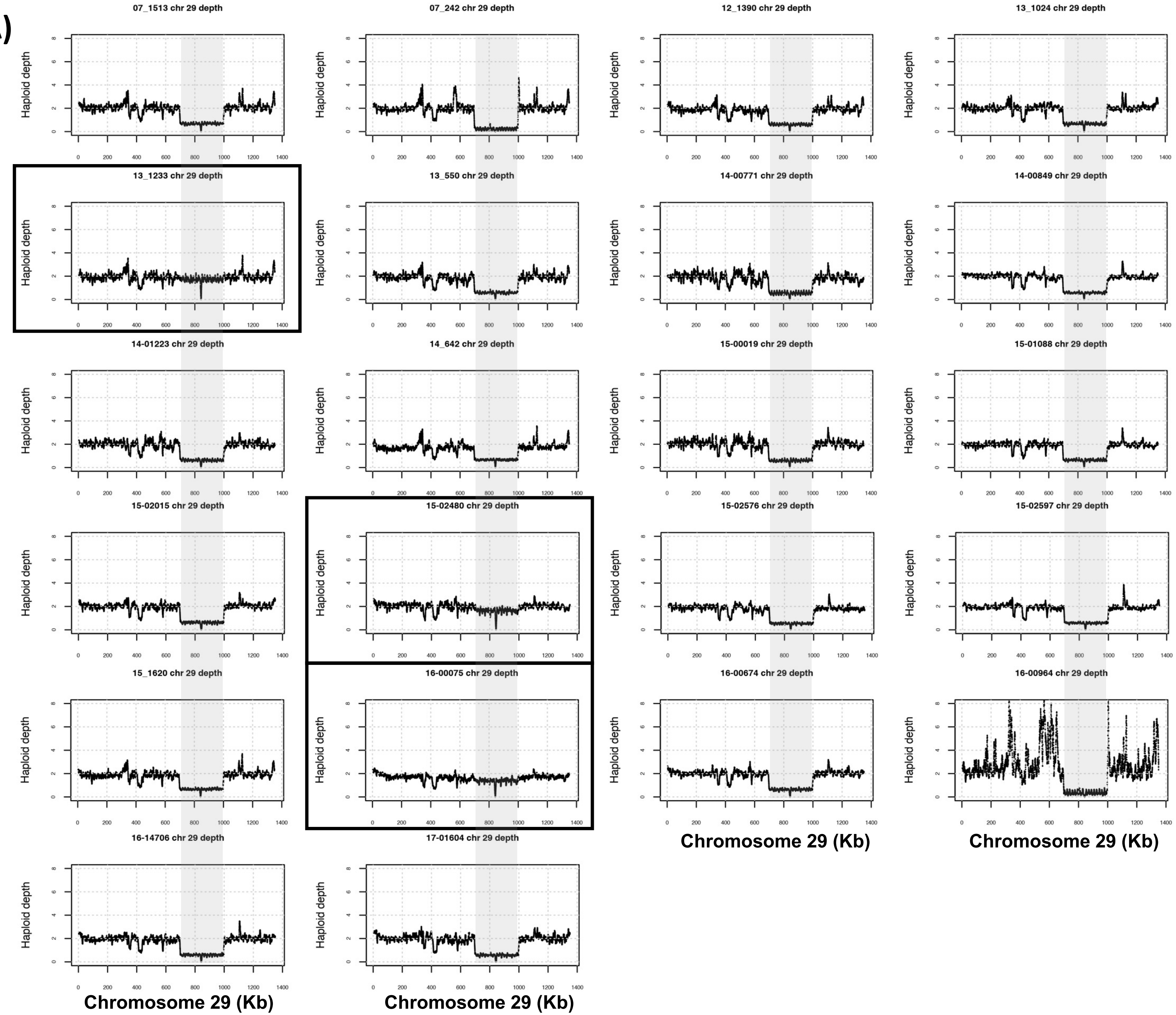


(B)

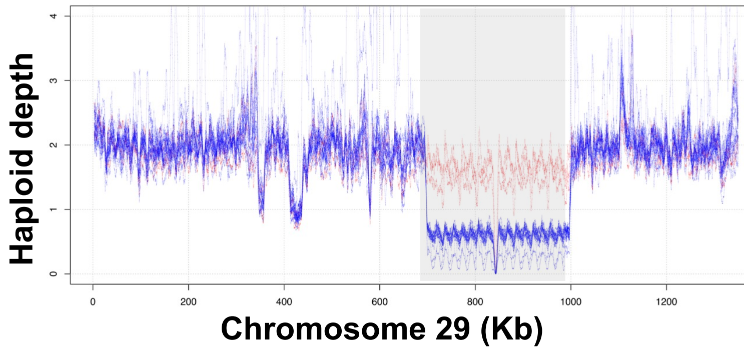


**Figure S8**. Chromosome 29 had a region at 704-1,006 Kb (302 Kb in length) with low depth (y-axis, normalised to haploid). (A) This region was significantly contracted in 19 isolates, but not the *L. tropica* reference genome and three genomes: 13_01233, 15_02480, 16_00075 (encircled by black boxes). (B) The depth of all 22 together is shown in the inset at the bottom contrasting the 19 (blue) vs three (red). All samples were disomic for chromosome 29.

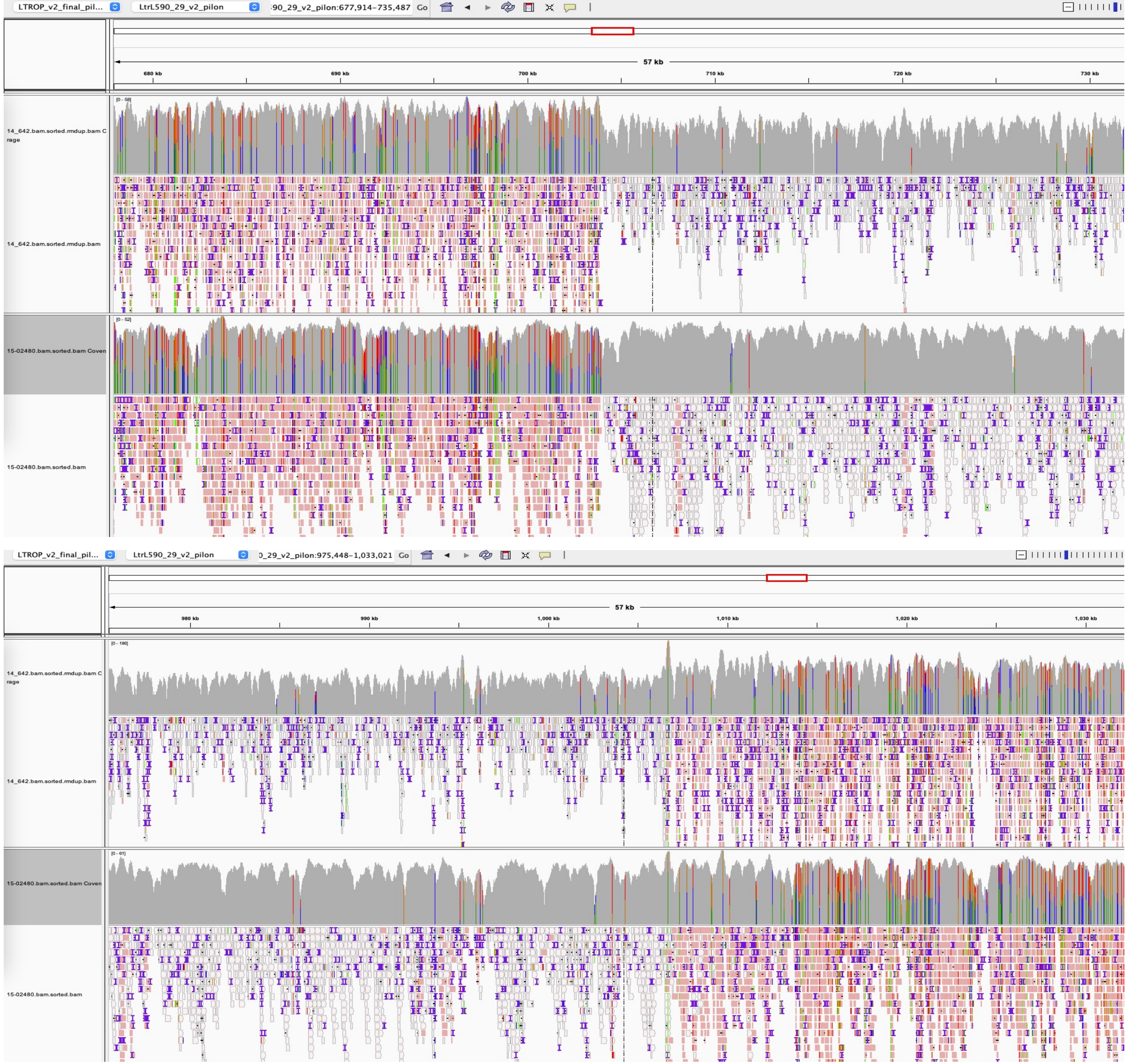


**Figure S9**. Visualisation of the reads mapped to the reference genome at chromosome 29 at 678-732 Kb (top) and at 977-1,031 Kb (bottom) showing no change in coverage in 15_020480 (middle) compared to 14_00642 (top) with the heterozygous deletion.

**Figure S10.** Assignment of genetically distinct population using FastBAPs for SNPs at chromosome 23. This shows chromosome 23's (A) 5' end (<250 Kb), (B) 3' end (>250 Kb) and (C) the whole chromosome showing the genetic relatedness of the 22 isolates with the *L. tropica* reference genome ("ref"). This shows unique ancestries at the 5' end with patterns related to the *L. tropica* reference in 07_00242, 13_01233, 15_02480, 16_00075 and 16_00964, contrasting with the 3' end and the overall genome-wide trend.

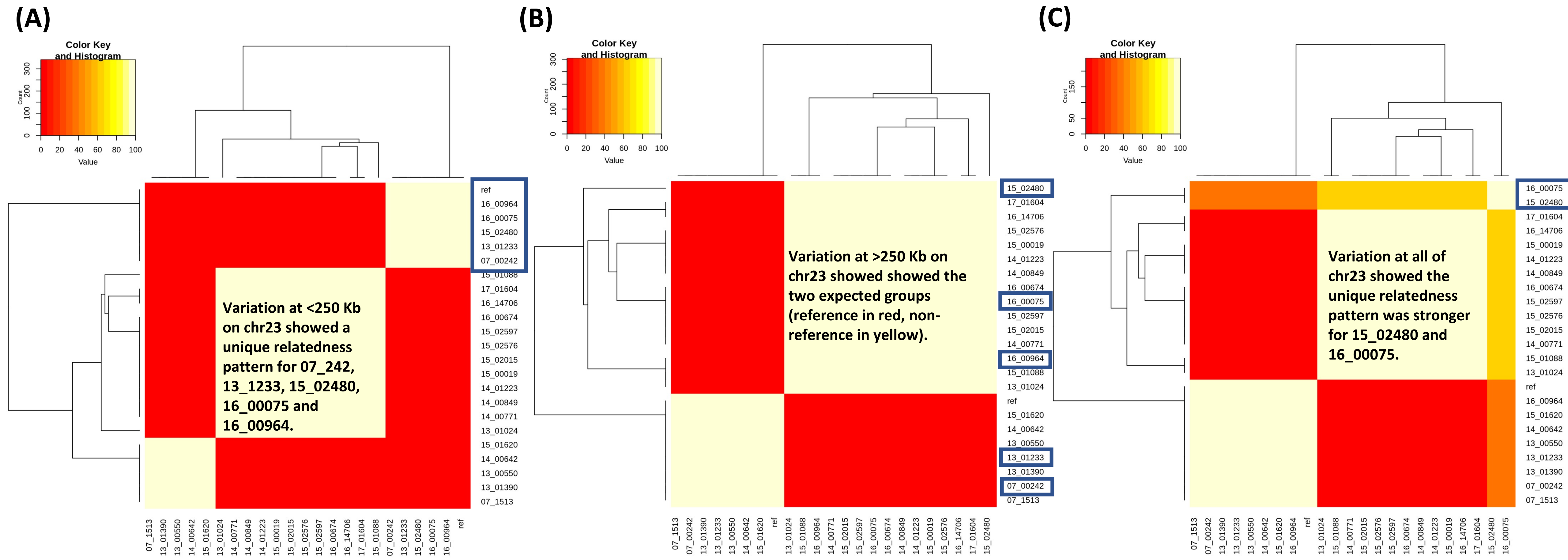

**Figure S11**. The read coverage and heterozygosity visualised for 14_00642's chromosome 10 reads for bases 0-340 Kb (top) and 240-580 Kb (bottom) using IGV. The top panel shows the log-scaled read coverage (grey) scaled from 0-244 (top) or 0-394 (bottom) with SNPs shown by colours. The bottom panel shows the reads and their SNPs. 14_642 had lower coverage at 20-250 Kb followed by higher coverage at <20 and 250 Kb. Chromosome 10's three potential strand-switch regions (SSRs) inferred from experiments in *L. major* are show by the red and blue panel below.

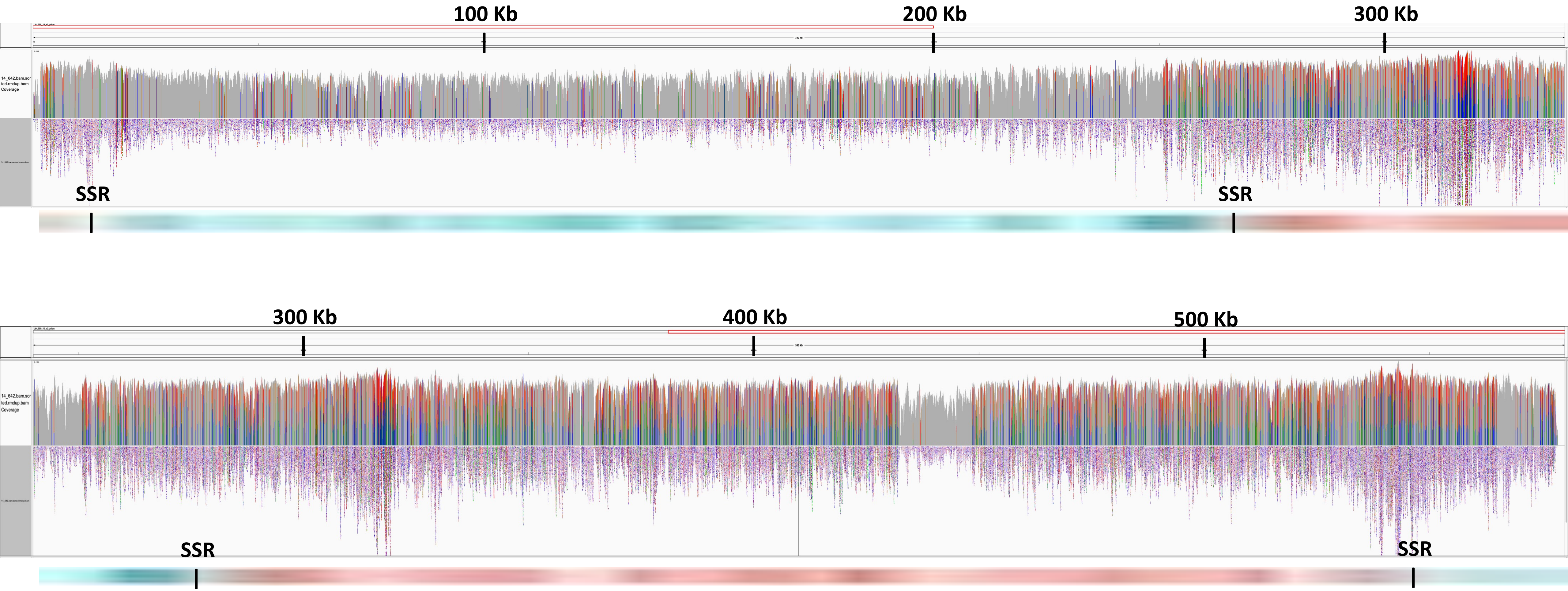

**Figure S12**. The read depth in isolates 15_02480 (top) and 14_00642 (bottom) when reads were mapped to the *L. tropica* reference genome (A) and then to their own *de novo* genome assemblies (B). (A) The median read depth of assemblies showed a similar pattern to the read-mapping to the reference genome. (B) The normalised haploid depth shows disomy for 15_02480, which was typical for all isolates bar 14_00642; and monosomy at 20-250 Kb followed by trisomy at <20 (grey shading) and 250 Kb (grey/yellow) for 14_00642. Chromosome 10 potentially has three strand-switch regions (SSRs) inferred from experiments in *L. major* (red and blue panel). The low level of heterozygous SNPs and high level of homozygous was shown by the numbers of valid SNPs across the read-depth allele frequency (RDAF) was not found in the other 20 isolates. (C) The homozygous (x-axis) and heterozygous (y-axis) SNPs per Kb for all 22 samples. 15_02480 and 14_00642 had much higher levels of homozygous SNPs on chromosome 10 compared to the other 20 isolates. (D) The RDAF distribution across the chromosome (green) showed shared mixed heterozygosity at 250-520 Kb (grey) with a change in heterozygosity symptomatic of an amplified region at >520 Kb (yellow).

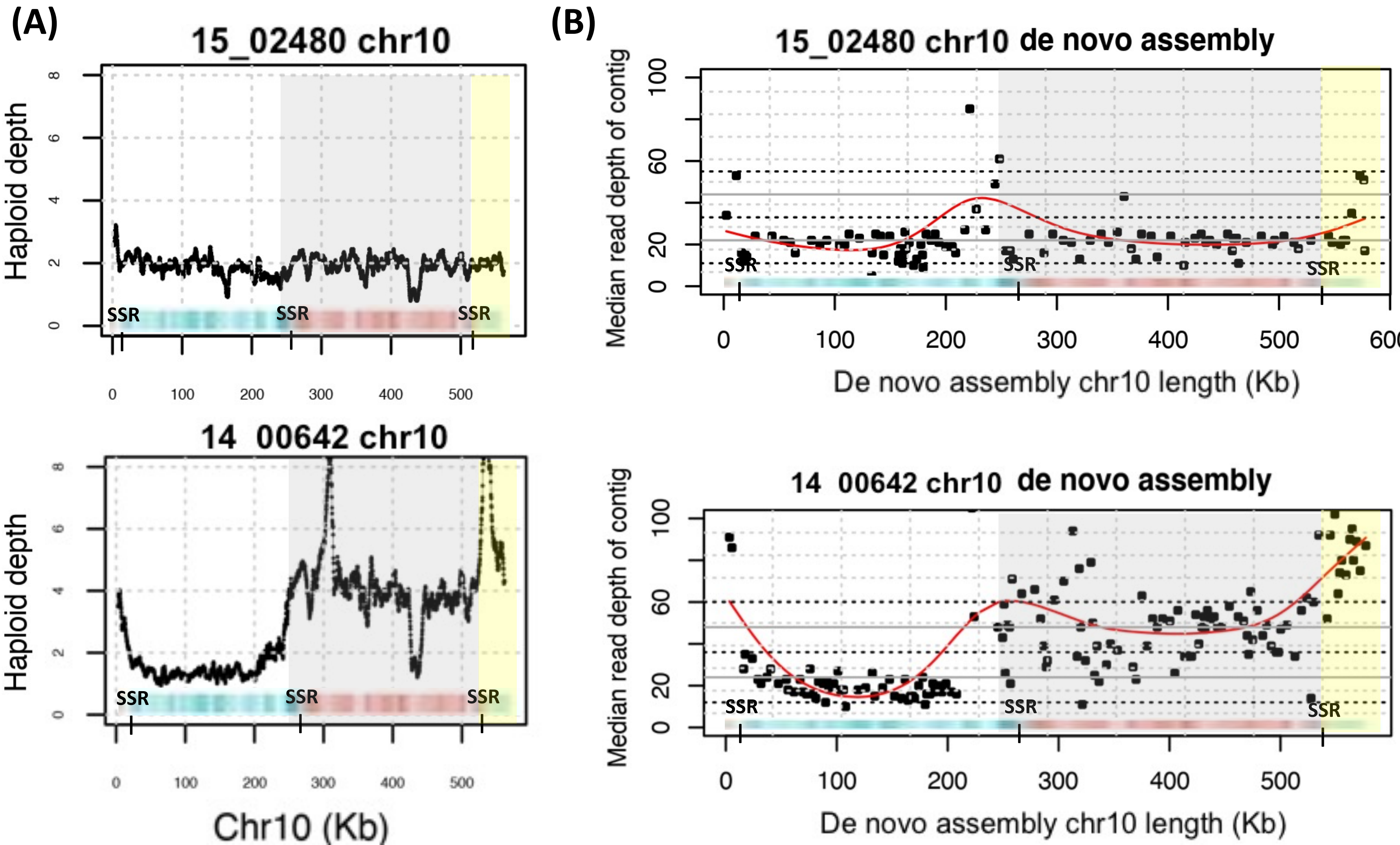


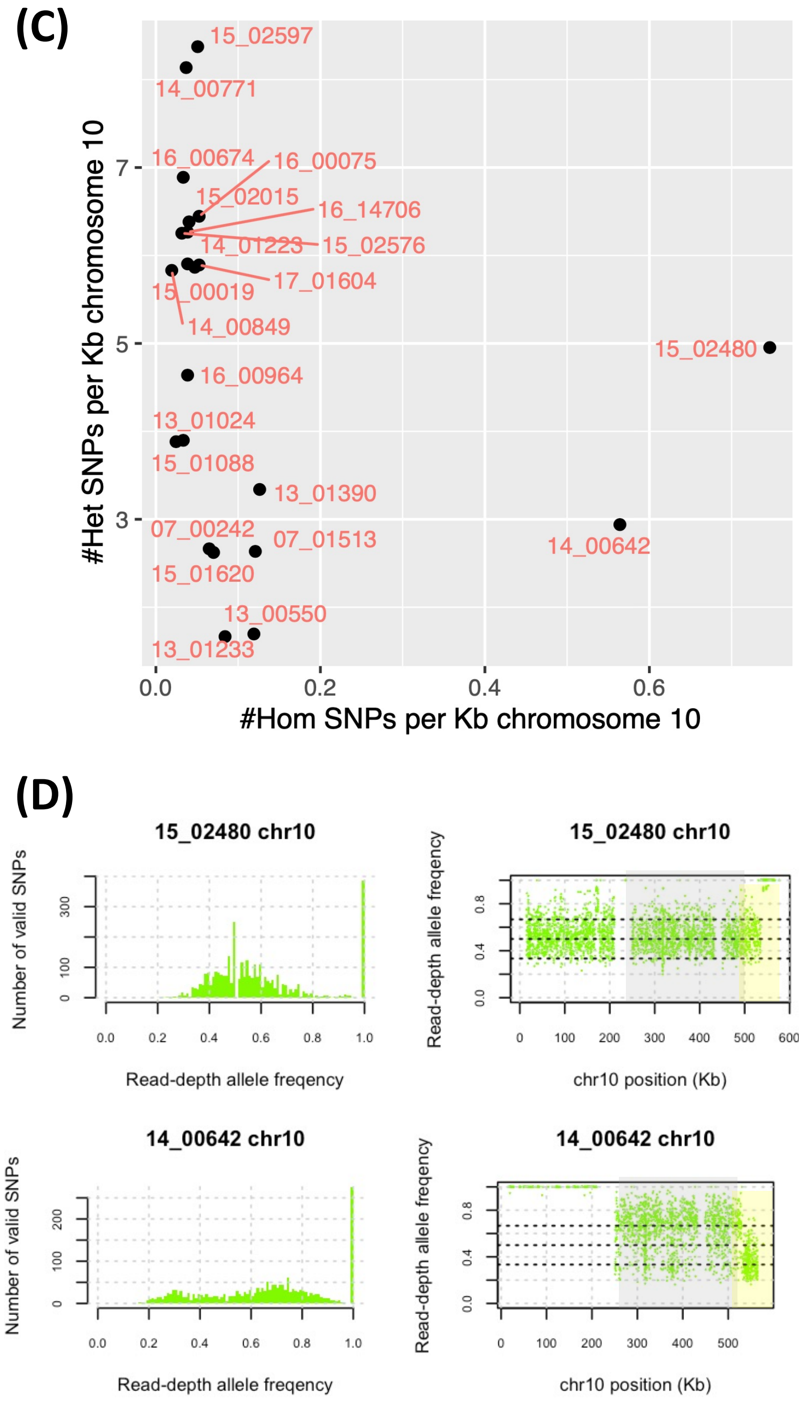

**Figure S13.** 07_00242 had homozygosity spanning the whole of chromosome 4 based on the read-depth allele frequency (RDAF) distribution (A) showing zero heterozygous SNPs and the RDAF across the chromosome showing minimal changes bar 720 homozygous SNPs. (B) A phylogeny constructed from chromosome 4's SNPs showing the relatedness of the 22 isolates with the *L. tropica* reference genome ("ref") showing that 07_00242 was genetically distinct. The inferred genetically distinct groups from FastBAPs are represented by the *L. tropica* reference (yellow area) and non-reference groups (purple area). (C) The homozygous (x-axis) & heterozygous (y-axis) SNPs per Kb for all 22 samples. There was no associations with *ori* regions / SSRs.

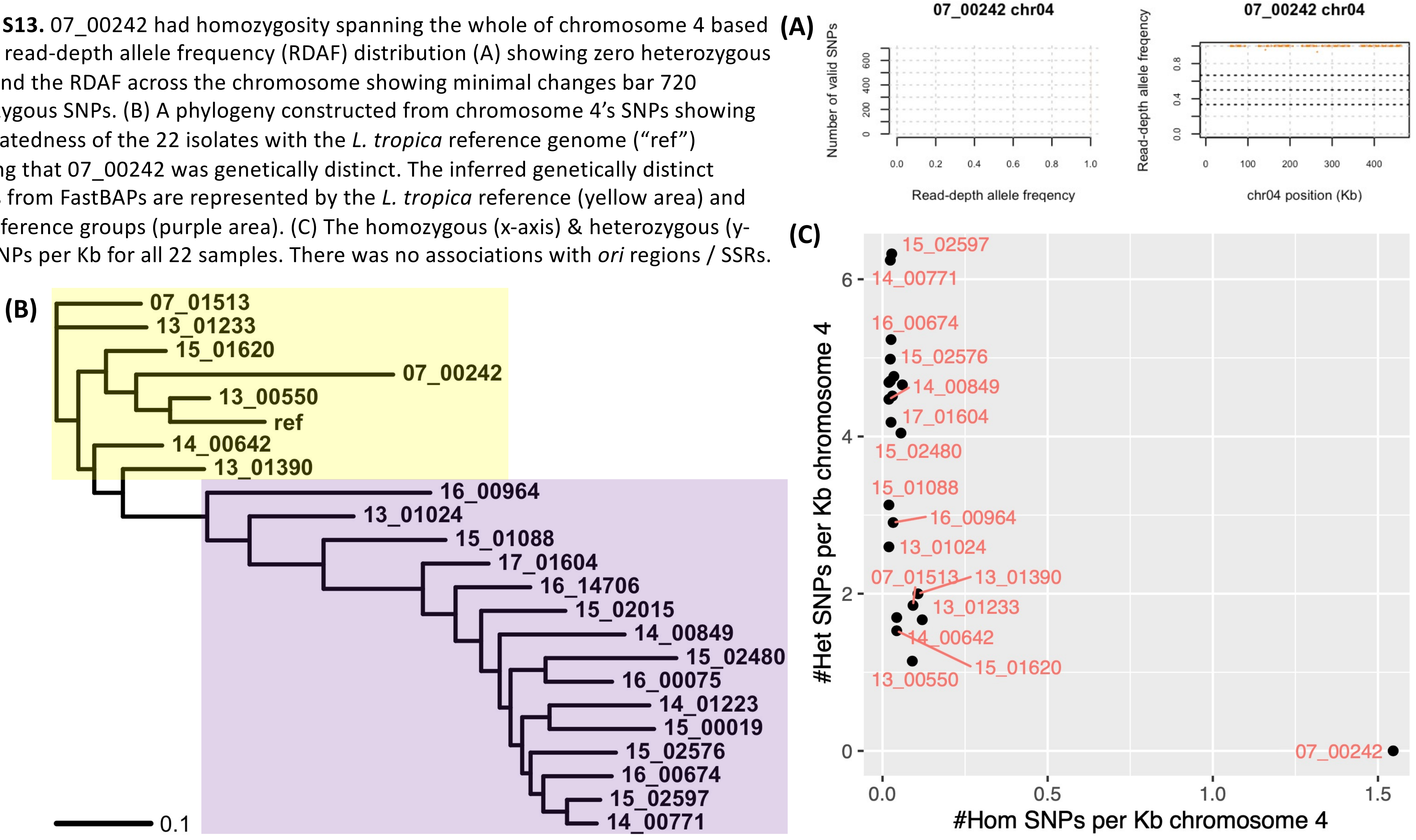

**Figure S14.** 07_01513 had evidence of heterozygosity followed by homozygosity >380 Kb on chromosome 11 where it had 652 homozygous (>7 times more than all the other samples). (A) This was illustrated by the SNPs' read-depth allele frequency (RDAF) distributions (left) and the RDAF levels across the chromosome (right). (B) The homozygous (x-axis) and heterozygous (y-axis) SNPs per Kb for all 22 samples. A putative *ori* region inferred from experiments in *L. major* is near 380 Kb. 07_01513 had intermediate di-/tri-somy at chromosome 11.

**(A)**

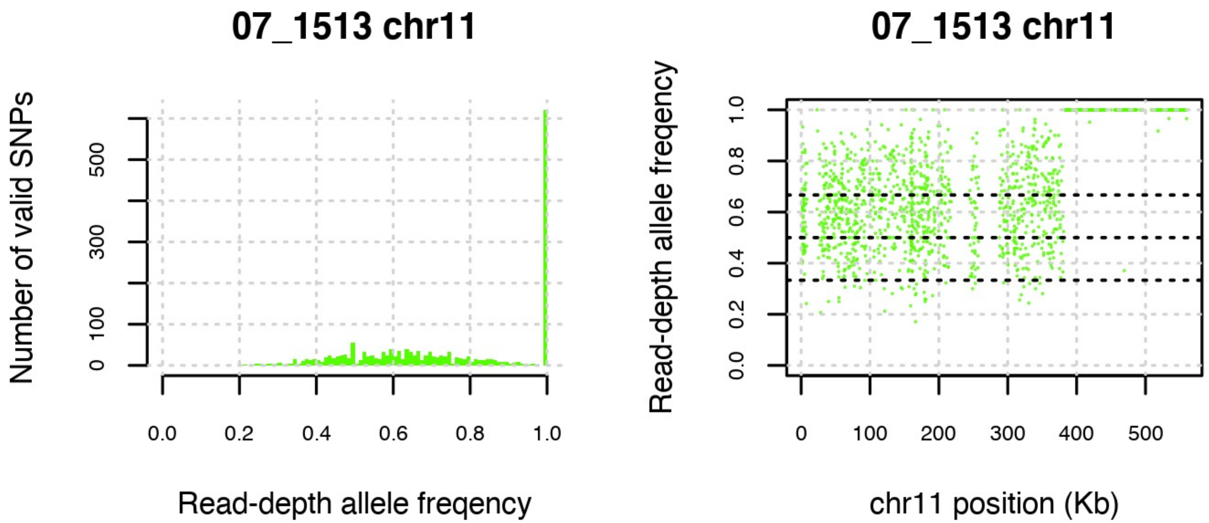


**(B)**

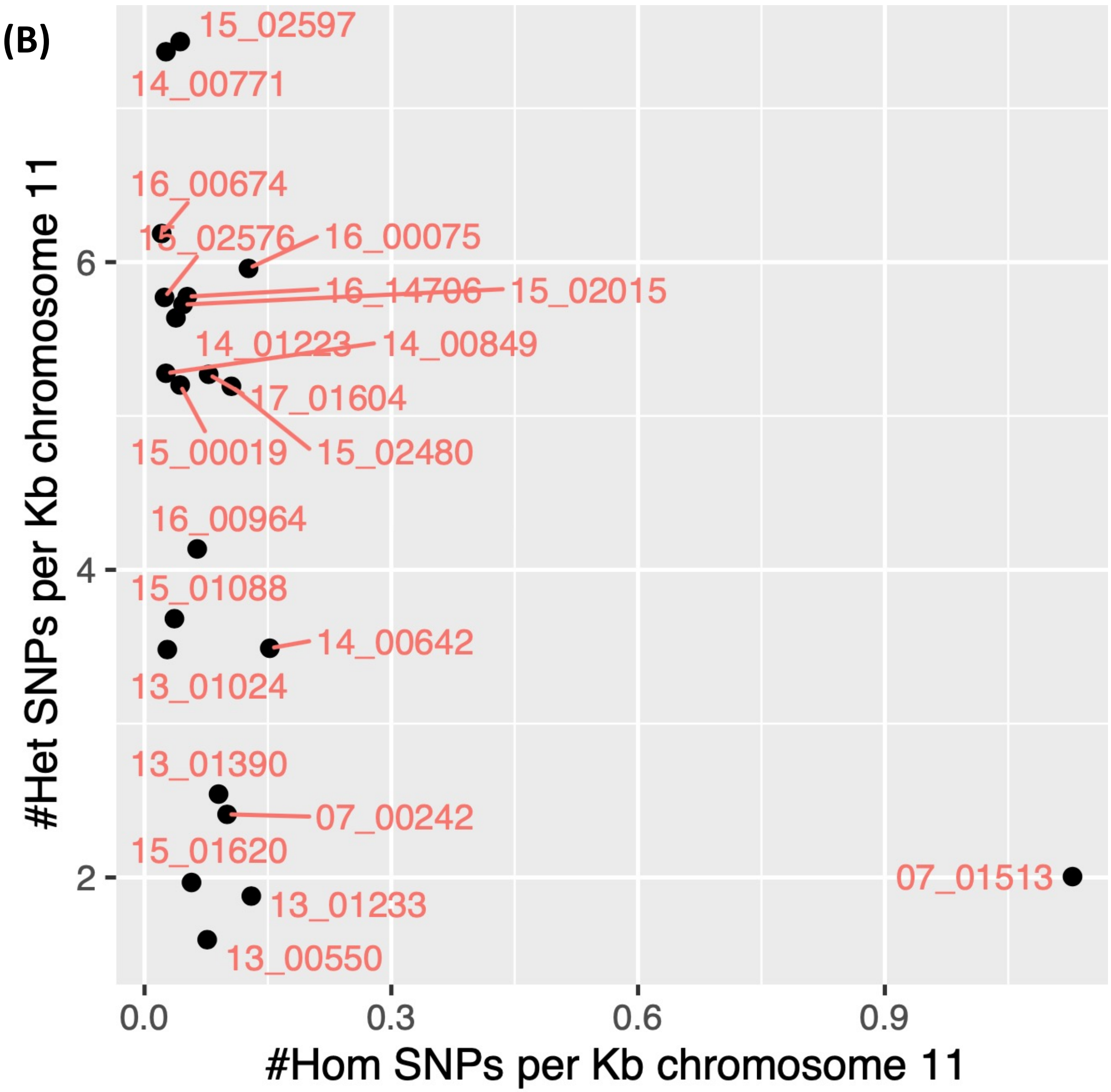

**Figure S15.** 15_02480 alone had a recombination breakpoint separating a heterozygous region at 0-330 on chromosome 12 from a homozygous one at >300 Kb. (A) This was illustrated by the read-depth allele frequency (RDAF) distribution across the chromosome mapped to the *L. tropica* reference genome (right) and the SNP frequency (left). (B) This region of homozygosity included a deletion at 400-450 Kb shown by the normalised read depth The other samples were disomic for chromosome 12. There were no associations with *ori* regions or SSRs. (C) The homozygous (x-axis) & heterozygous (y-axis) SNPs per Kb for all 22 samples. (D) The haploid depth for 15_02480's reads mapped to its *de novo* genome assembly suggesting a novel region at 400-450 Kb. (E) IGV visualisation of the reads mapped to the reference genome at chromosome 12 at 392-462 Kb showing the deletion in 14_00642 (top) vs 15_02480 (bottom).

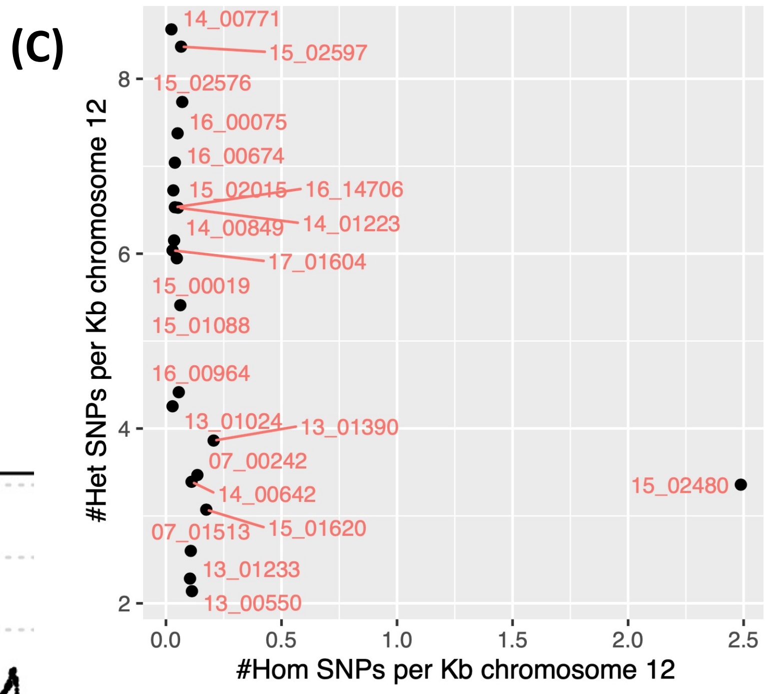


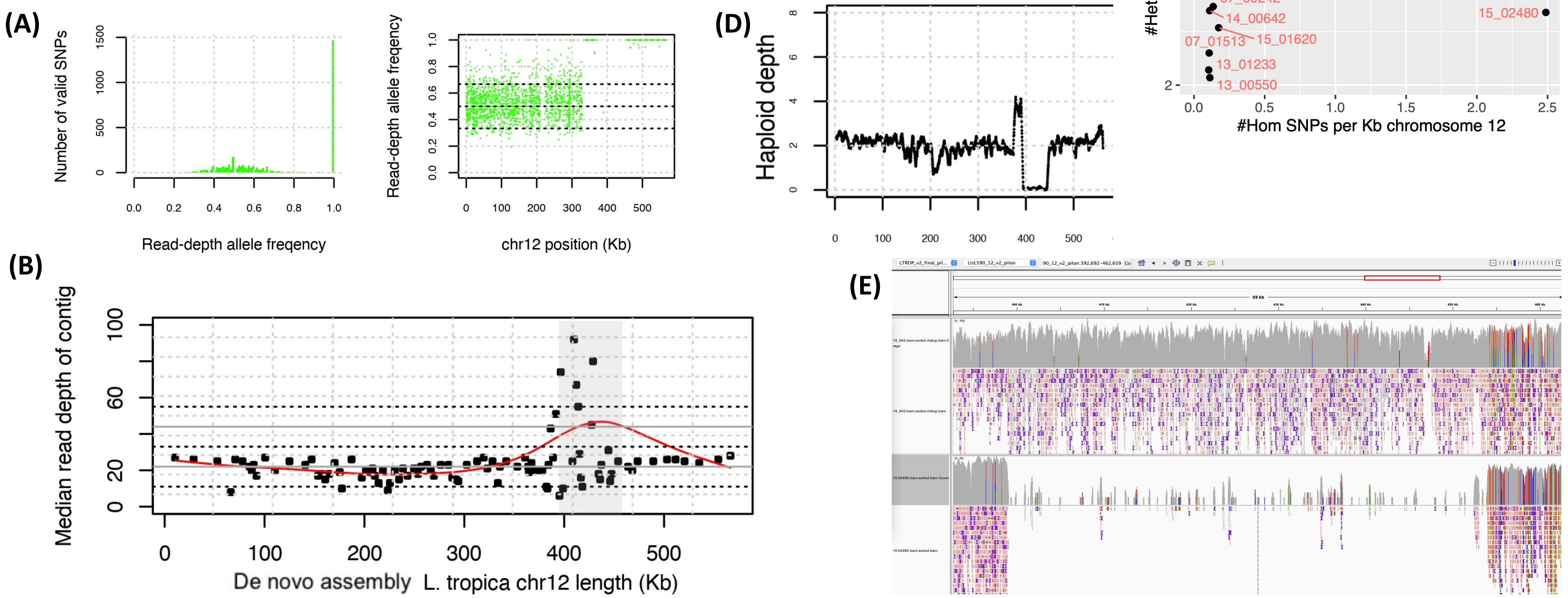

**Figure S16.** 07_00242 and 16_00964 had evidence of recombination breakpoints separating a homozygous region at >530 Kb on chromosome 13 from heterozygous regions 5′ of this. (A) This was illustrated by the SNPs' RDAF distributions (left) and the RDAF levels across the chromosome (right). (B) A phylogeny constructed from chromosome 13's SNPs showing the relatedness of the 22 isolates with the *L. tropica* reference genome ("ref") showing that 07_00242 and 16_00964 are somewhat genetically distinct. The inferred genetically distinct groups from FastBAPS are represented by the *L. tropica* reference (yellow area) and non-reference groups (purple area): 16_00964's variation at this chromosome was assigned to the reference group. (C) The homozygous (x-axis) & heterozygous (y-axis) SNPs per Kb for all 22 samples. 07_242 (somy 2.24) and 16-00964 (somy 1.89) were about disomic for chromosome 13. There were no associations with *ori* regions or SSRs.

**(A)**

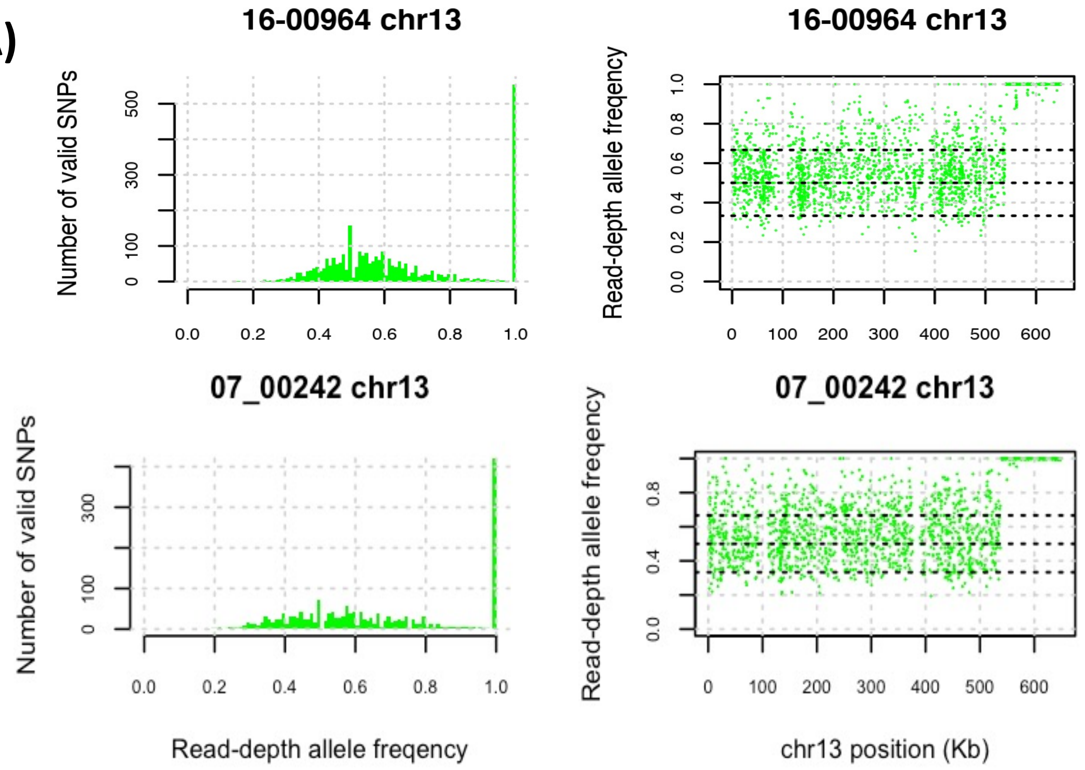


**(B)**

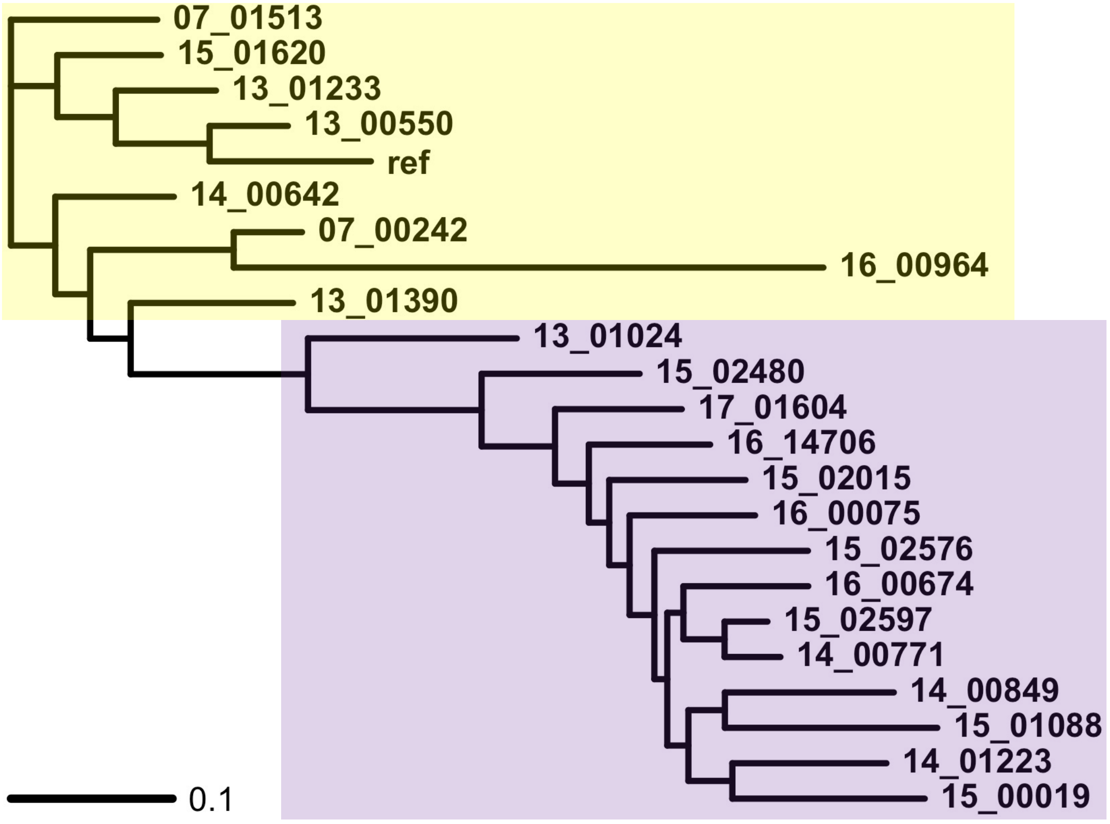


**(C)**

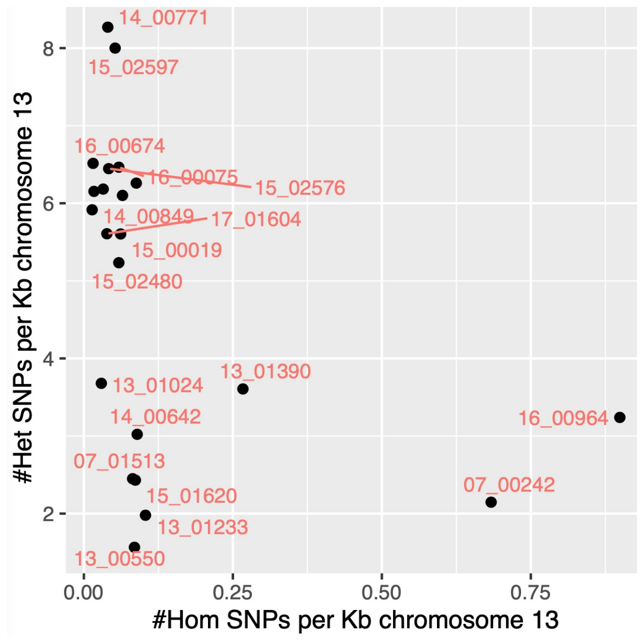

**Figure S17.** 14_00642 had evidence of homozygosity at <70 Kb on chromosome 14 where it had 213 homozygous SNPs across the chromosome (90% more all the other samples), followed by heterozygosity >70 Kb. (A) This was illustrated by the SNPs' read-depth allele frequency (RDAF) distributions (left) and the RDAF levels across the chromosome (right). (B) The homozygous (x-axis) and heterozygous (y-axis) SNPs per Kb for all 22 samples. 14_00642 was tri-/tetra-somic at chromosome 14 (somy 3.55). There were no associations with *ori* regions or SSRs. The lengths of the *de novo* assembled chromosome 14 excluding N bases were similar to that of the reference (638.2±24.6 vs 651.0 Kb)

**(A)**

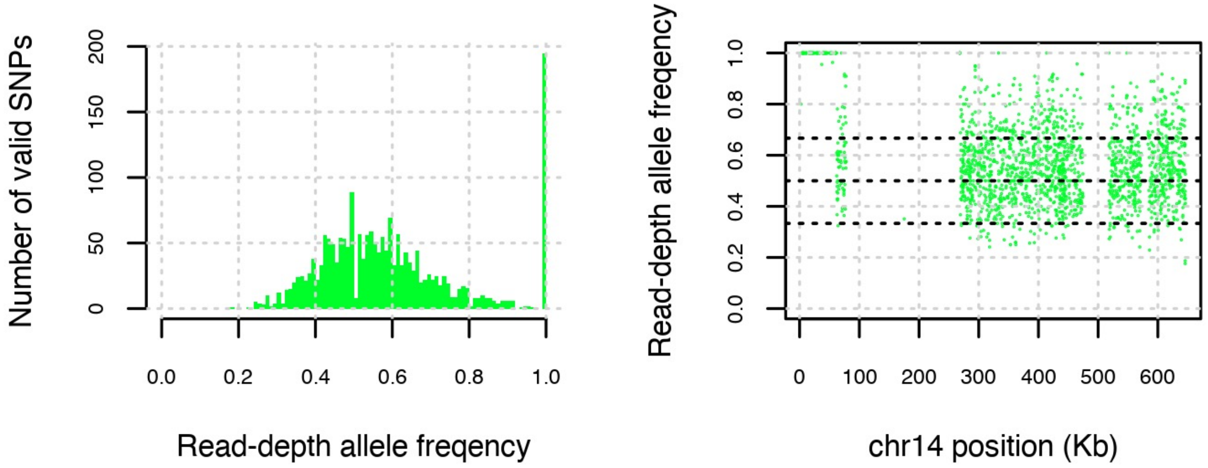


**(B)**

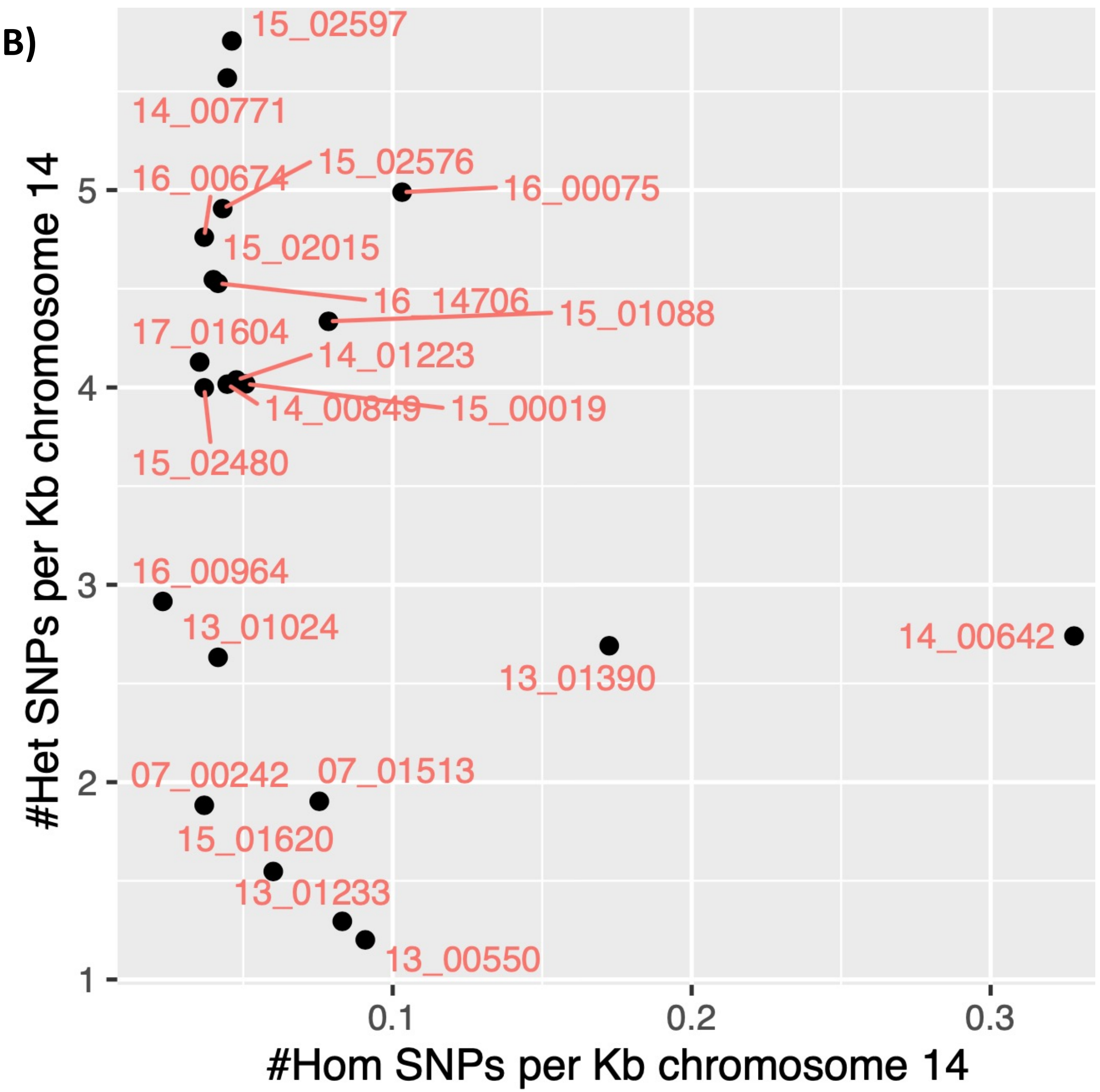

**Figure S18.** 07_00242 had homozygosity at <80 Kb on chromosome 17 following by heterozygosity. (A) This is based on the read-depth allele frequency (RDAF) distribution (left) and across the chromosome (right). This sample had 189 homozygous SNPs across the chromosome (>66% more than any other sample). (B) The homozygous (x-axis) and heterozygous (y-axis) SNPs per Kb for all 22 samples. Most samples including 07_00242 were approximately disomic for chromosome 17. There were no associations with *ori* regions or SSRs.

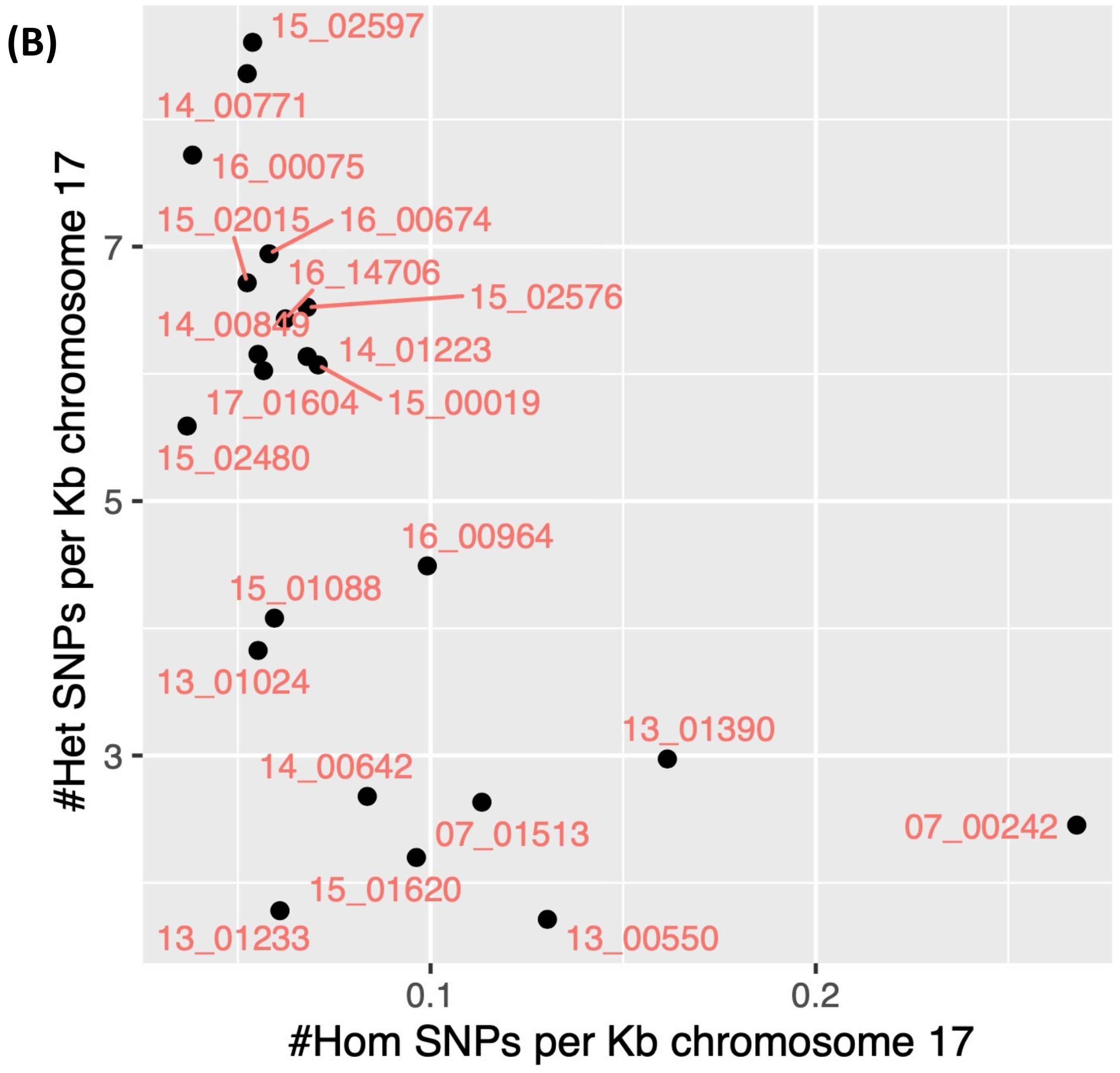


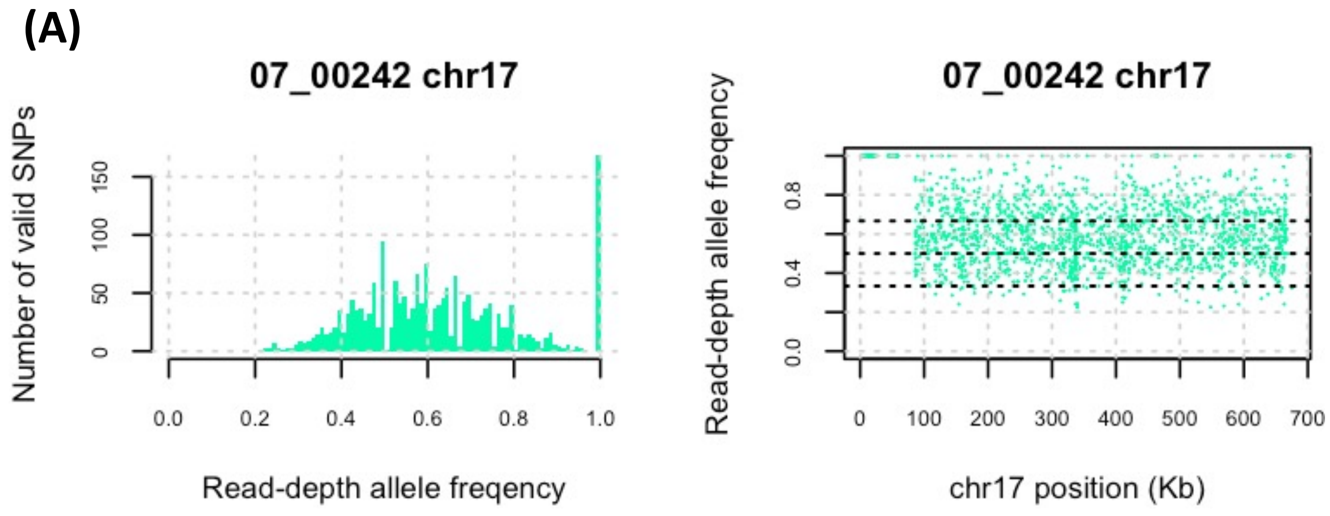

**Figure S19.** 07_00242 had homozygosity for all of chromosome 22. (A) This was based on the read-depth allele frequency (RDAF) distribution (left) and the dearth of heterozygous SNPs (right). (B) 14_00642 had a homozygous region at <90 Kb followed by a region with mixed heterozygosity for the remainder. (C) The homozygous (x-axis) and heterozygous (y-axis) SNPs per Kb for all 22 samples. Like most samples, 07_00242 was disomic for this chromosome, whereas 14_00642 had an intermediate level between di- and tri-somy. There were no associations with *ori* regions or SSRs.

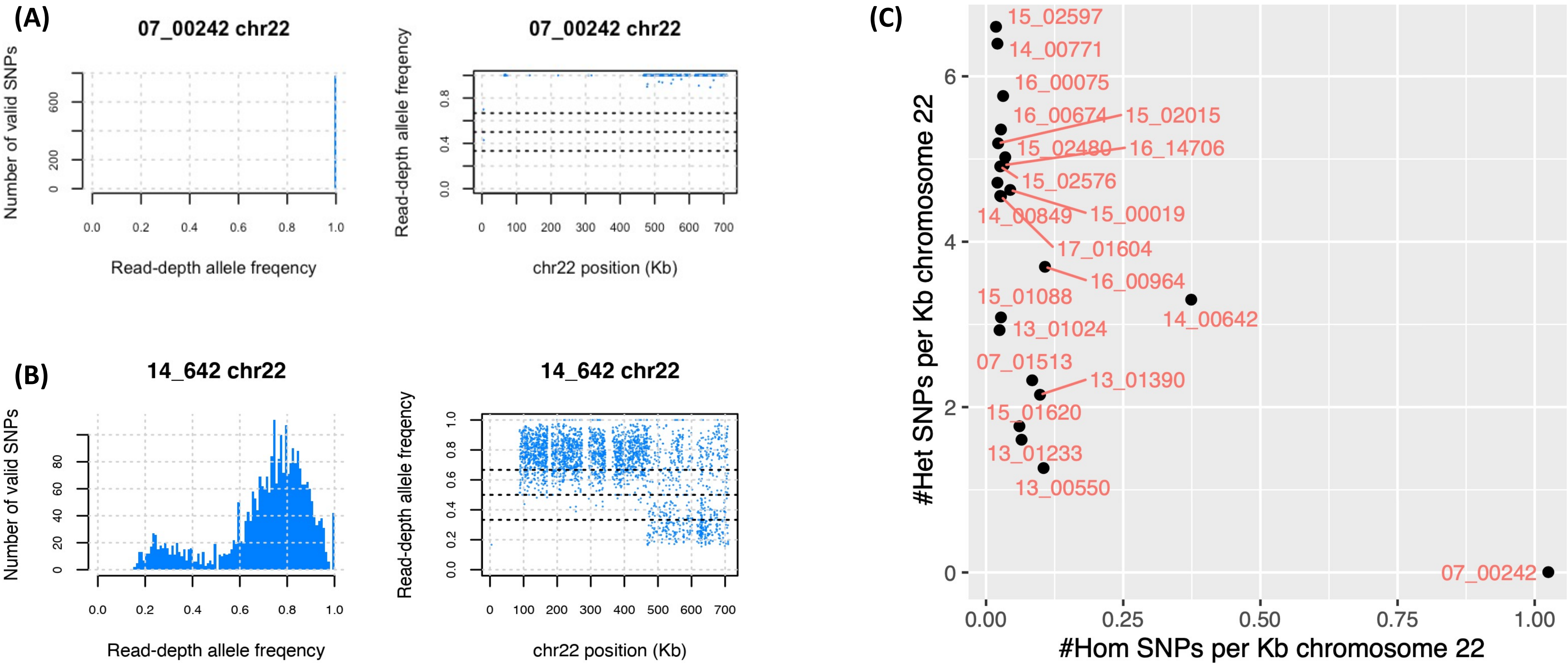

**Figure S20.** 07_01513 (A) and 14_00771 (B) had evidence of recombination breakpoints separating a homozygous region at <90 Kb in 14_00771 and <150 Kb in 07_01513 on chromosome 24 from heterozygous regions 3' of this. 14_00771 also had a region with a higher average read-depth allele frequency (RDAF) at 90-200. This was illustrated by the SNPs' RDAF distributions (left) and the RDAF levels across the chromosome (right). (C) The homozygous (x-axis) and heterozygous (y-axis) SNPs per Kb for all 22 samples. All 22 samples were approximately disomic for chromosome 24. There were no associations with *ori* regions or SSRs.

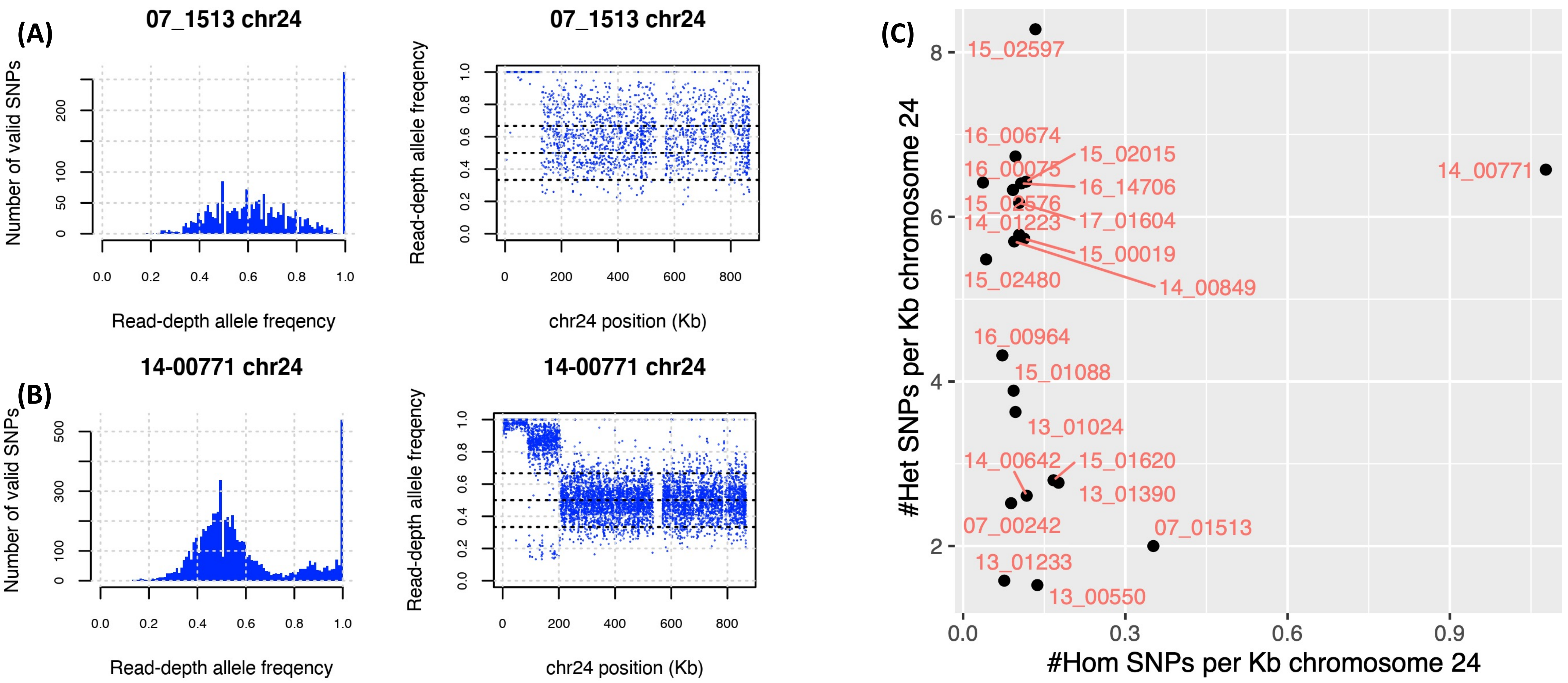

**Figure S21.** 13_01390 had evidence of quasi-homozygosity on chromosome 27 where it had 1,626 homozygous (>8 times more than all the other samples). (A) This was illustrated by the SNPs' read-depth allele frequency (RDAF) distributions (left) and the RDAF levels across the chromosome (right). (B) A phylogeny constructed from chromosome 27's SNPs showing the relatedness of the 22 isolates with the *L. tropica* reference genome ("ref") showing that 13_01390 was somewhat genetically distinct. The inferred genetically distinct groups from FastBAPs are represented by the *L. tropica* reference (yellow area) and non-reference groups (purple area). (C) The homozygous (x-axis) & heterozygous (y-axis) SNPs per Kb for all 22 samples. A putative *ori* region inferred from experiments in *L. major* is near 1.05 Mb separating a more heterozygous region 3' of the centromere. 13_01390 had near disomy at chromosome 27 (somy 1.77).

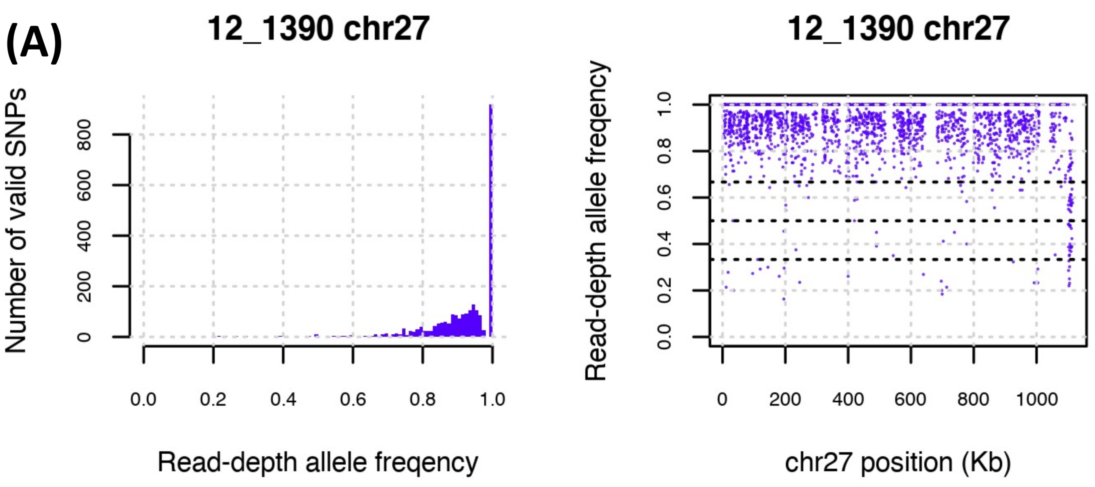


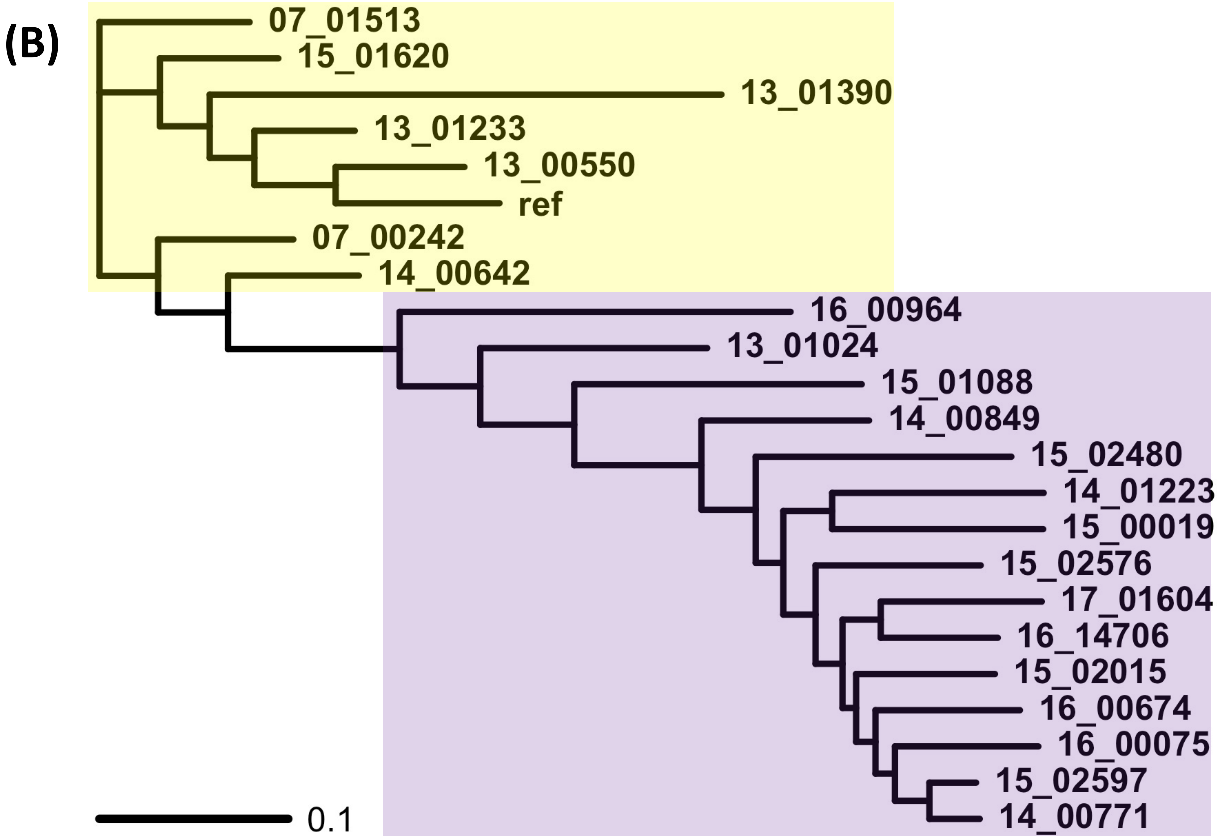


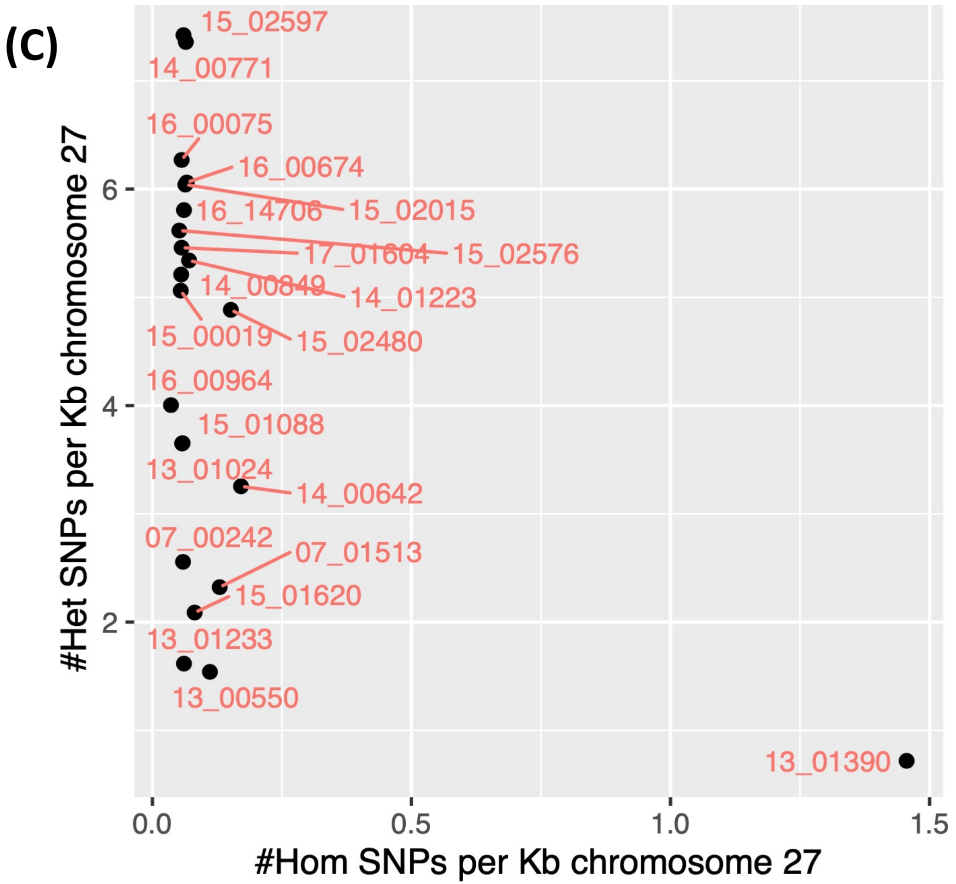

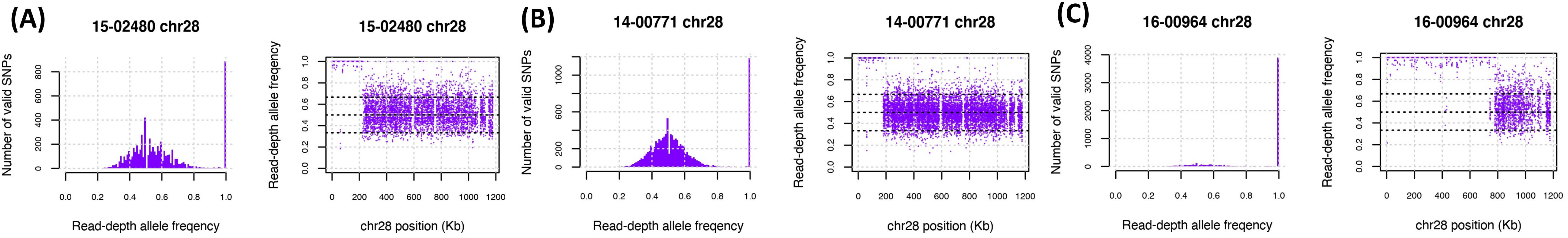


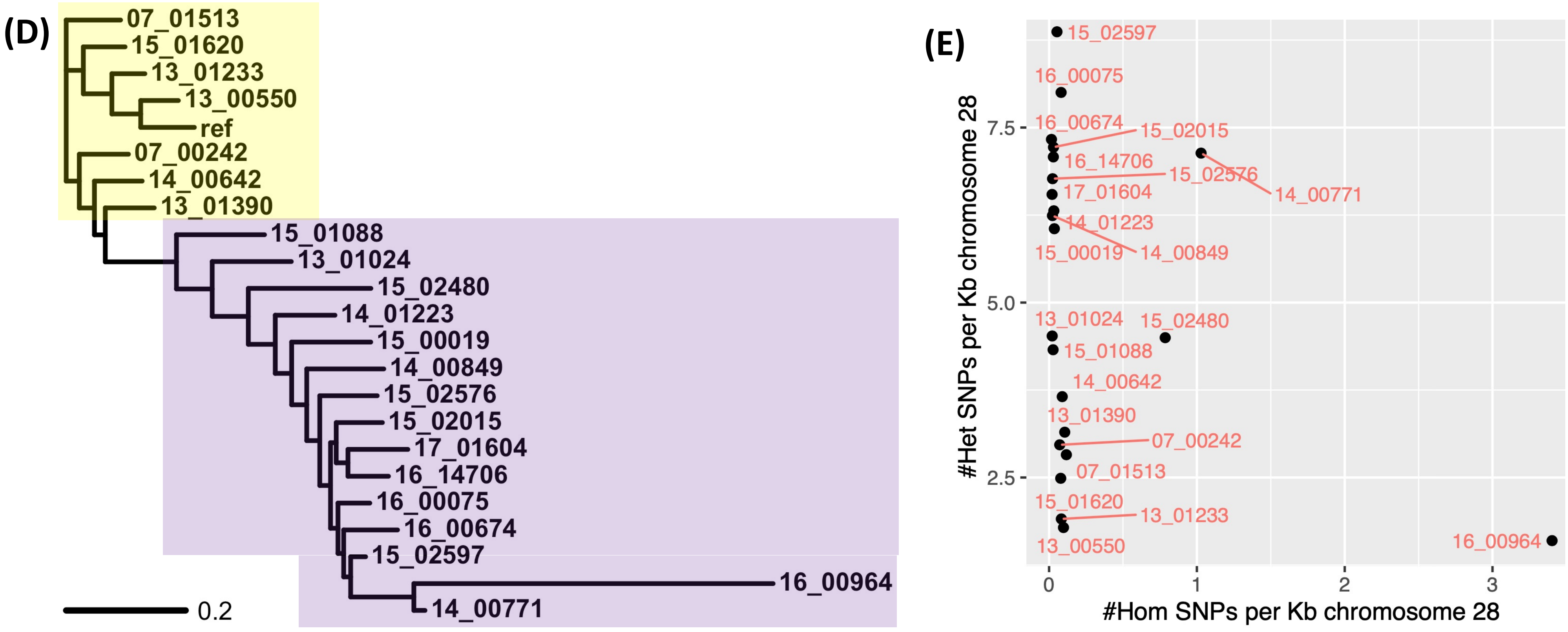


**Figure S22.** 15_02480 (A), 14_00771 (B) and 16_00964 (C) had evidence of recombination breakpoints separating a homozygous region at <180 Kb in 14_00771, at <210 Kb in 15_02480, and at <740 Kb in 16-00964 on chromosome 28 from heterozygous regions 3' of this. This was illustrated by the SNPs' RDAF distributions (left) and the RDAF levels across the chromosome (right). All 22 samples were approximately disomic for chromosome 28. There were no associations with *ori* regions or SSRs. (D) A phylogeny constructed from chromosome 28's SNPs showing the relatedness of the 22 isolates with the *L. tropica* reference genome ("ref") showing that 16_00964 was genetically distinct with some SNPs shared with 14_00771. The inferred genetically distinct groups from FastBAPs are represented by the *L. tropica* reference (yellow area) and non-reference groups (purple area). (E) The homozygous (x-axis) and heterozygous (y-axis) SNPs per Kb for all 22 samples.

**Figure S23.** 15_02015 had evidence of a homozygous region at <120 Kb on chromosome 30 where it had 479 homozygous (more than twice any other sample). (A) This was illustrated by the SNPs' read-depth allele frequency (RDAF) distributions (left) and the RDAF levels across the chromosome (right). (B) The homozygous (x-axis) and heterozygous (y-axis) SNPs per Kb for all 22 samples. Most samples including 15_02015 were approximately disomic for chromosome 30. There were no associations with *ori* regions or SSRs. The length of the *de novo* chromosome 30 assembly for 15_01088 (corrected p=0.009) was shorter than the reference genome's chromosome 30.

(A)

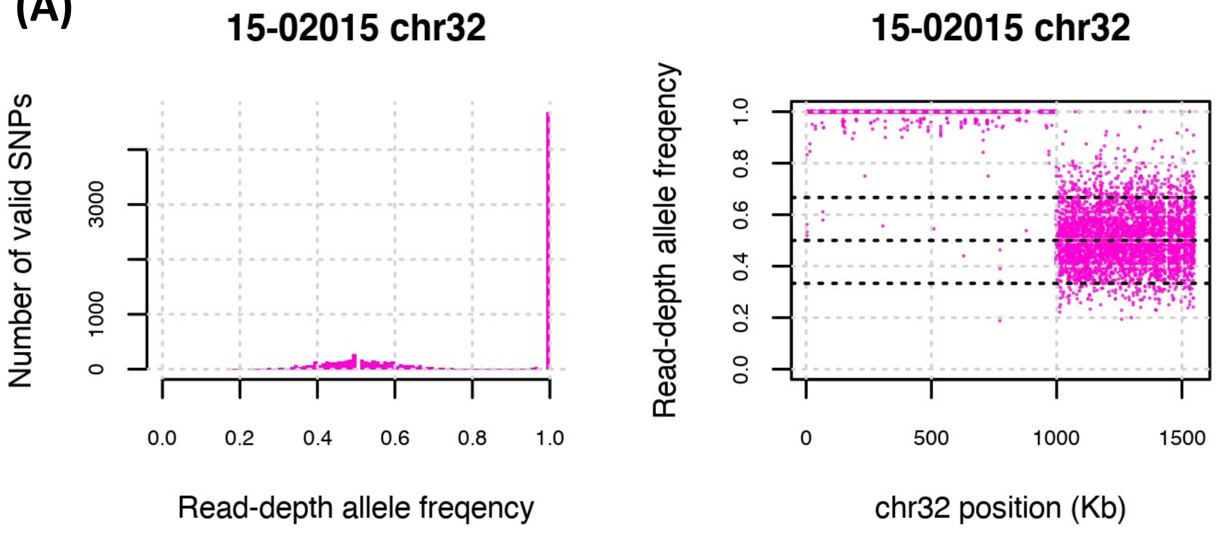


(B)

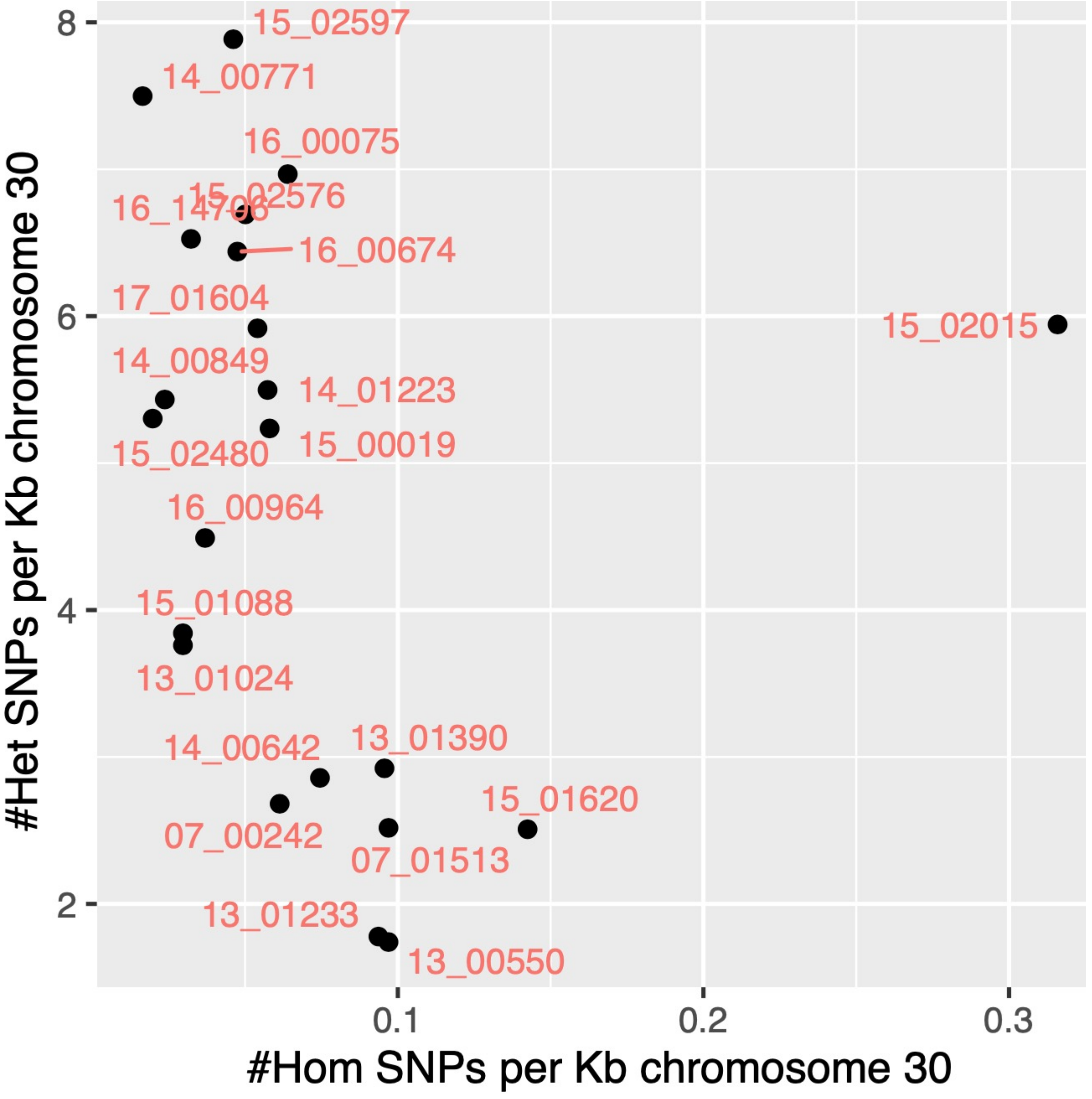

**Figure S24.** 07_00242 had heterozygosity at <1.2 Mb on chromosome 33 following by mainly homozygosity >1.2 Mb. (A) This was based on the read-depth allele frequency (RDAF) distribution (left) showing 3,537 heterozygous SNPs and the RDAF across the chromosome 1,363 homozygous SNPs. (B) The homozygous (x-axis) and heterozygous (y-axis) SNPs per Kb for all 22 samples. There were no associations with *ori* regions or SSRs. Chromosome 33 had a mix of somy levels across samples.

(B)

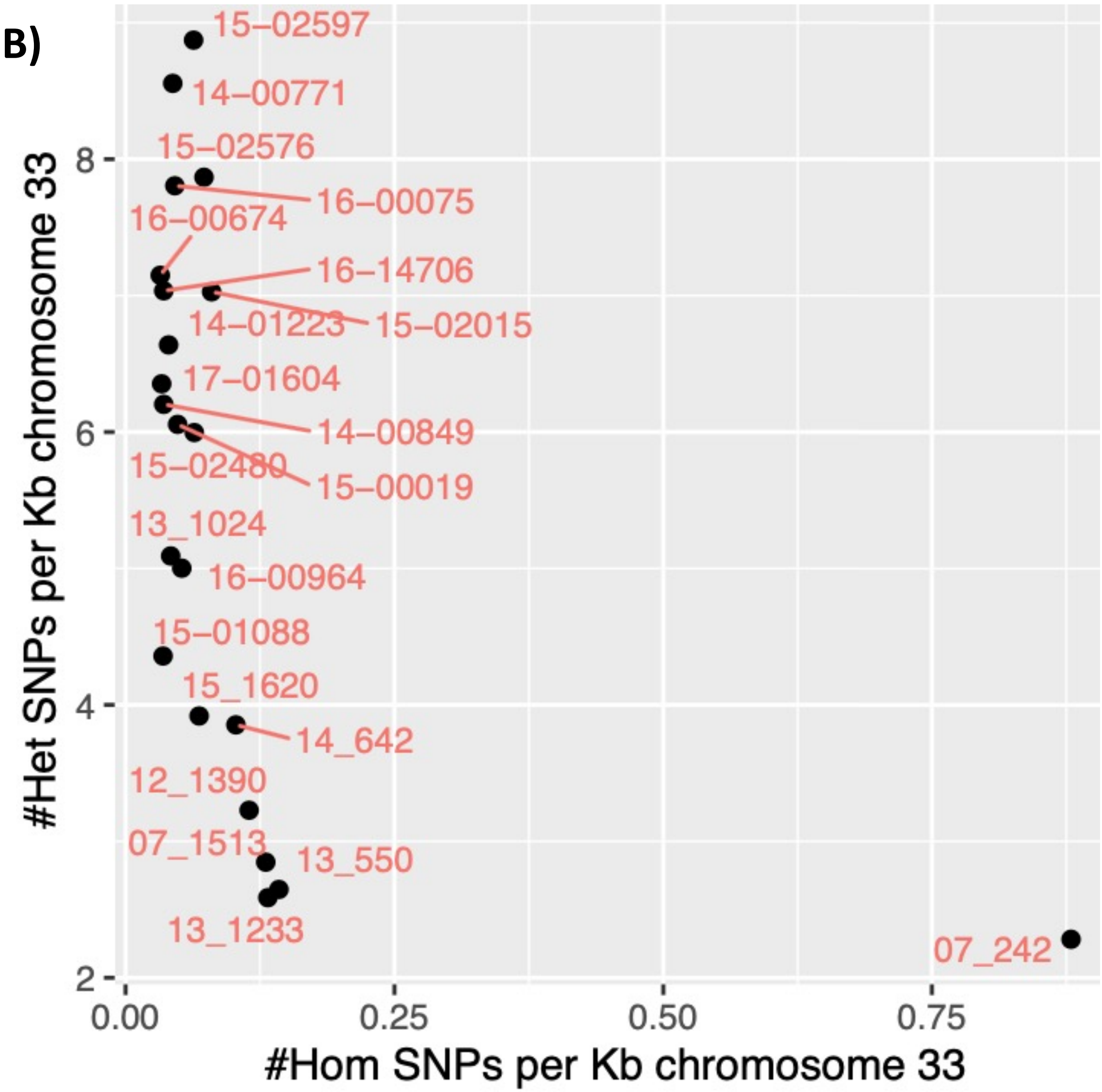


(A)

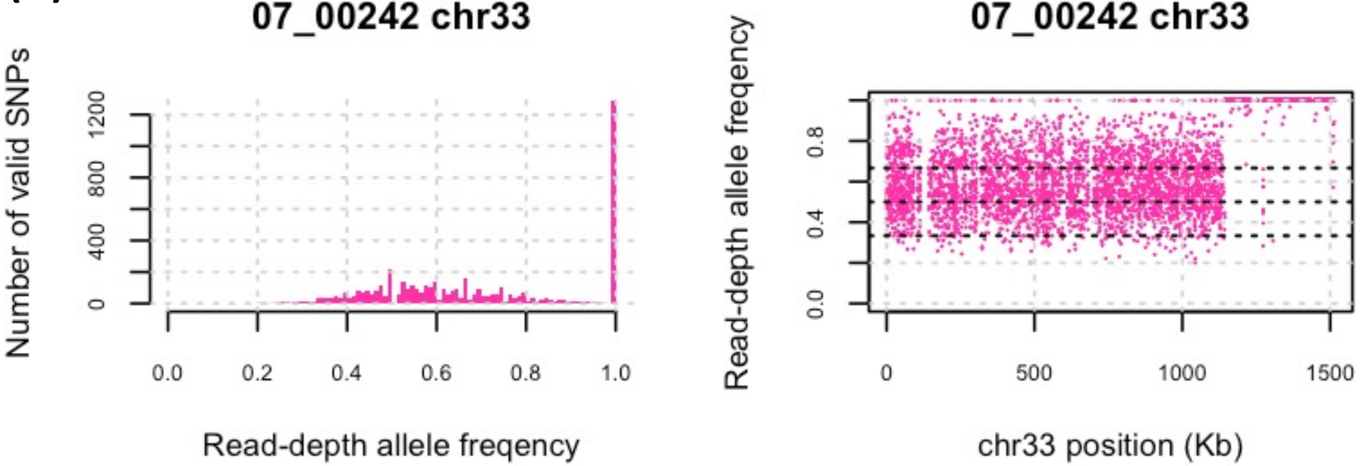

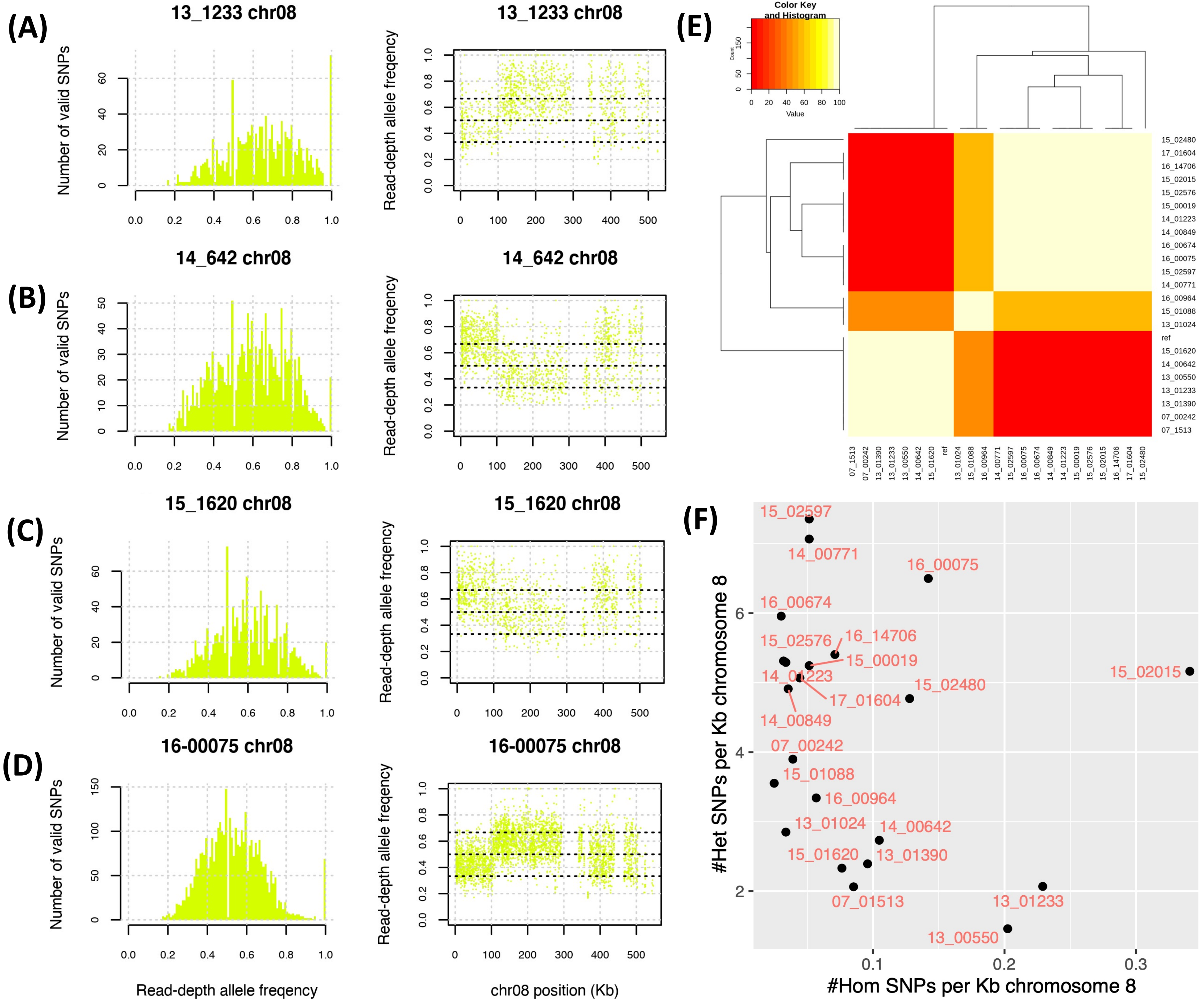


**Figure S25.** There was evidence of recombination breakpoints separating regions with a lower average read-depth allele frequency (RDAF) at 110-380 Kb from one with higher average RDAF at <110 Kb and >380 Kb on chromosome 8. This was associated with intermediate di-/tri-somy in 13_01233 (A) (somy 2.56), 14_00642 (B) (somy 2.50), 15_01620 (C) (somy 2.35) and 16-00075 (D) (somy 2.66). This was illustrated by the SNPs' RDAF distributions and the RDAF levels across the chromosome. (E) Assignment of genetically distinct population using FastBAPs for all SNPs at chromosome 8 showing the genetic relatedness of the 22 isolates with the *L. tropica* reference genome ("ref"), indicating an intermediate classification for 13_01024, 15_01088 and 16_00964. (F) The homozygous (x-axis) and heterozygous (y-axis) SNPs per Kb for all 22 samples. The other samples were approximately disomic for chromosome 8 (somy range 1.65-2.25) except 07_00242 (somy 3.50). There were no associations with *ori* regions or SSRs. The length of the *de novo* chromosome 8 assembly for 15_01088 (p=0.028) was shorter than the reference genome's chromosome 8.

**Figure S26.** 15_02597 (A) and 14_00771 (B) and had numerous recombination breakpoints separating short regions of homozygosity on chromosome 25 from heterozygous regions. This was illustrated by the SNPs' RDAF distributions (left) and the RDAF levels across the chromosome (right). (C) The homozygous (x-axis) and heterozygous (y-axis) SNPs per Kb for all 22 samples. Most samples including these were approximately disomic for chromosome 25. There were no associations with *ori* regions or SSRs.

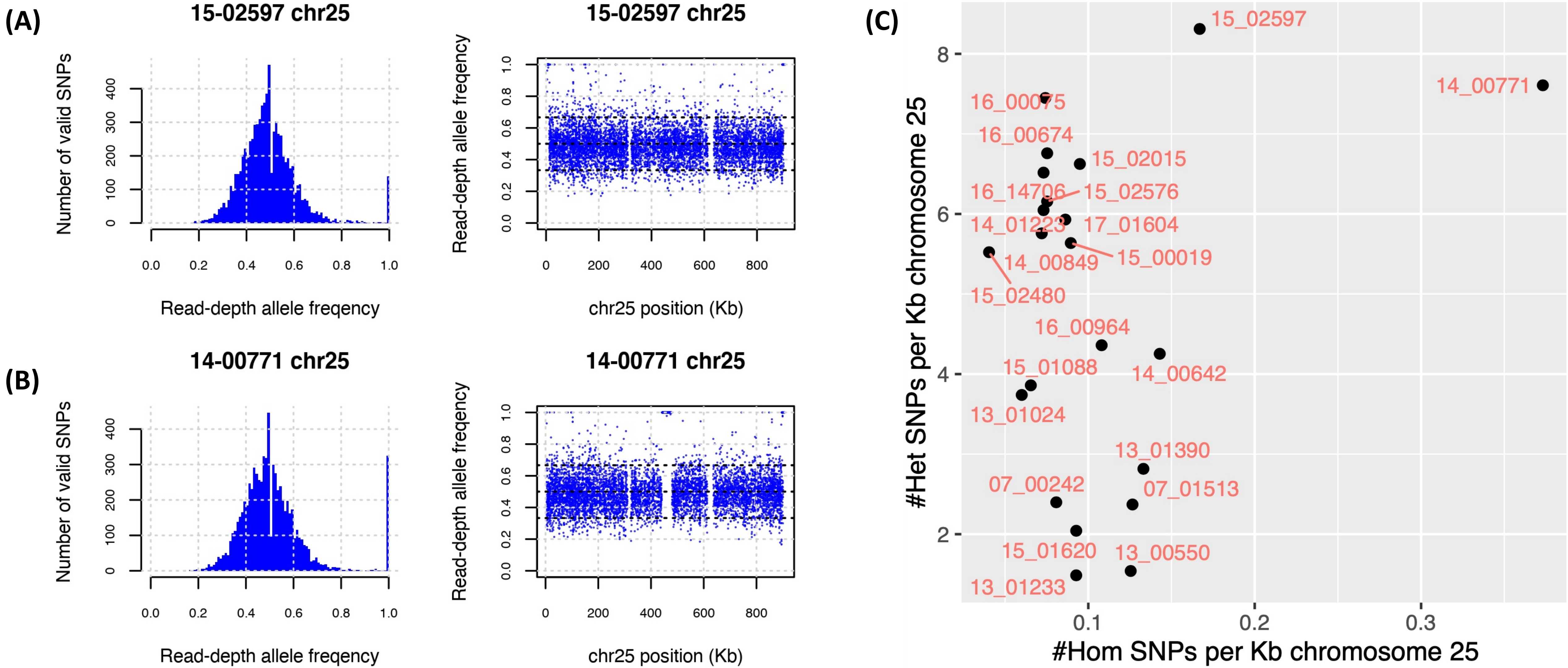

**Figure S27.** 15_02015 (A) 07_00242 (B) and had regions of homozygosity on chromosome 32 spanning the whole chromosome for 07_00242 based on the read-depth allele frequency (RDAF) distribution (left) and the dearth of heterozygous SNPs (right). 15_02015 had a homozygous region at <1.0 Mb followed by a heterozygous region for the remainder. There were no associations with *ori* regions or SSRs. Most samples including these were approximately disomic for chromosome 32. (C) A phylogeny constructed from chromosome 32’s SNPs showing the relatedness of the 22 isolates with the *L. tropica* reference genome (“ref”) showing that 07_00242 and 15_02015 were genetically distinct. The inferred genetically distinct groups from FastBAPs are represented by the *L. tropica* reference (yellow area) and non-reference groups (purple area). (D) The homozygous (x-axis) and heterozygous (y-axis) SNPs per Kb for all 22 samples. The lengths of the *de novo* chromosome 32 assemblies for 07_242 (p=0.046) and 14_00642 (p<0.001) were longer than the reference genome’s chromosome 32.

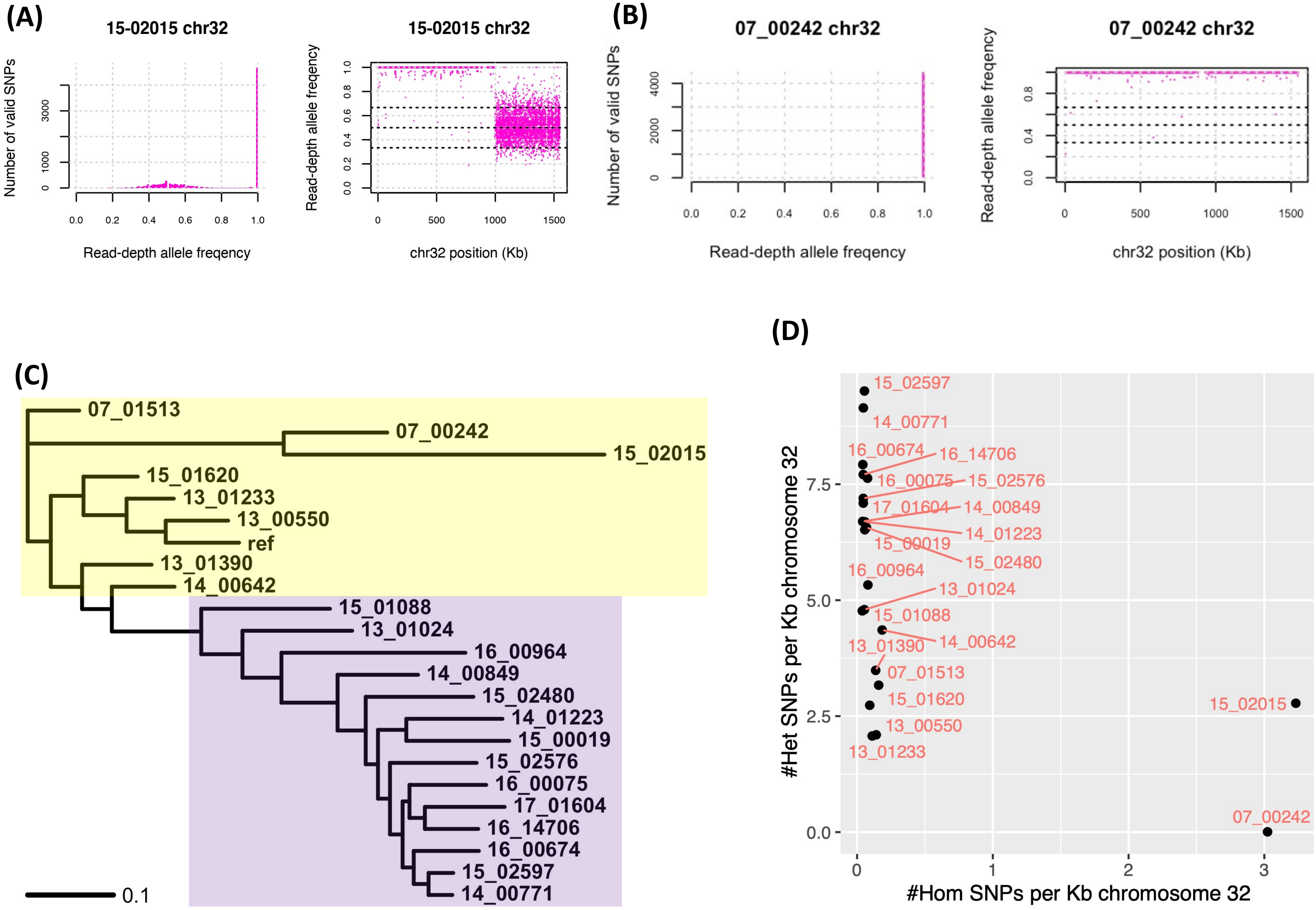

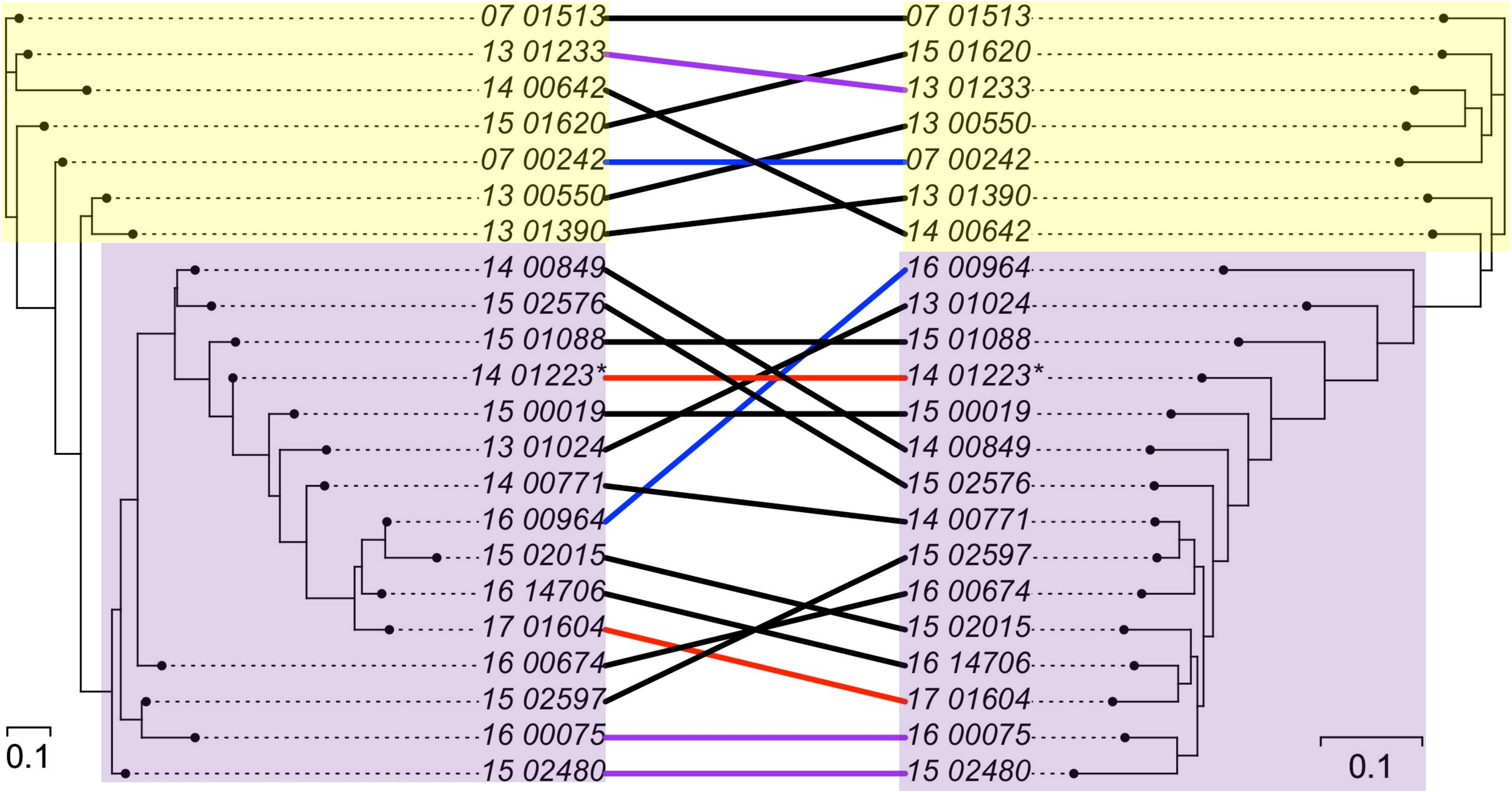


**Figure S28**. A co-phylogeny constructed from kDNA (left) and genome-wide (right) SNPs showing the relatedness of the 22 isolates. The samples were from Syria (n=17, black or red connections), Afghanistan (n=3, purple connections) and Iran (n=2, blue connections). Isolates 14_01223 and 17_01604 were from the same patient (red connections). The inferred genetically distinct groups from FastBAPs are represented by the reference-related group (yellow area) and non-reference group (purple area).
